# Distinct activation mechanisms underlie ligand efficacy at a GPCR

**DOI:** 10.64898/2026.10.12.758633

**Authors:** Matthew K. Howard, Eve J. Fine, Karthik Srinivasan, Daniel D. Richman, Jerome Freudenberg, Zara Weinberg, Jingyou Rao, Ziyue Zou, Christian Macdonald, Patrick Rockefeller Grimes, Balazs R. Varga, James S. Fraser, Mark Von Zastrow, Susruta Majumdar, Harold Pimentel, Justin English, Ron O. Dror, Aashish Manglik, Willow Coyote-Maestas

**Affiliations:** Department of Pharmaceutical Chemistry, UCSF, San Francisco, CA, USA; Department of Bioengineering and Therapeutic Science, UCSF, San Francisco, CA, USA; TETRAD graduate program, UCSF, San Francisco, CA, USA; Department of Bioengineering, Stanford University, Stanford, CA, USA; Department of Computer Science, Stanford University, Stanford, CA, USA; Department of Molecular and Cellular Physiology and Structural Biology, Stanford University School of Medicine, Stanford, CA, USA; Institute for Computational and Mathematical Engineering, Stanford University, Stanford, CA, USA; Bioinformatics Interdepartmental Program, UCLA, Los Angeles, CA, USA; Center for Clinical Pharmacology, Department of Anesthesiology and Washington University Pain Center, Washington University School of Medicine, Saint Louis, MO, USA; Department of Psychiatry and Behavioral Sciences, University of California, San Francisco, San Francisco CA, USA; Department of Cellular and Molecular Pharmacology, University of California, San Francisco, San Francisco, CA, USA; Quantitative Biology Institute, University of California, San Francisco, San Francisco, CA, US; Department of Computer Science, UCLA, Los Angeles, CA, USA; Department of Computational Medicine, David Geffen School of Medicine, UCLA, Los Angeles, CA, USA; Department of Human Genetics, David Geffen School of Medicine, UCLA, Los Angeles, CA, USA; Department of Biochemistry, University of Utah School of Medicine, Salt Lake City, UT, USA; Department of Anesthesia and Perioperative Care, University of California, San Francisco, CA, USA; Chan Zuckerberg Biohub, San Francisco, CA, 94158, USA

## Abstract

Ligands that bind to the same G protein-coupled receptor (GPCR) can produce widely different levels of signaling. These differences in efficacy are central to pharmacology, yet how a receptor sequence determines signaling level remains incompletely understood. Here we develop a deep mutational pharmacology approach to measure how each residue in the µ-opioid receptor (µOR) shapes responses to 15 opioids, yielding ∼400,000 measurements of variant effects on signaling and expression, including full concentration-response series for three agonists. Mutational effects on potency are strongest near the orthosteric pocket, whereas effects on efficacy extend through the transmembrane core and intracellular face. Ligands share many functional determinants, but sensitivity to mutation differs with efficacy. As efficacy decreases, the residues required for signaling shift from the binding pocket towards the intracellular face; partial agonists therefore depend on residues that are dispensable for full agonists. Mutations can broadly increase receptor responses, allowing antagonists such as naloxone to produce signaling comparable to a full agonist and increasing the efficacy of partial agonist ligands. Combined mutations can also reorder the relative efficacies of ligands, showing that sequence changes reshape pharmacological differences rather than simply increasing or decreasing all responses together. These findings support a model in which ligands of different efficacy activate the receptor through distinct allosteric mechanisms which are determined by residues distributed across the receptor. Mapping these contributions across diverse ligands with deep mutational pharmacology provides a way to understand how one receptor converts molecular recognition into a range of signaling outputs.

## Introduction

G protein-coupled receptors (GPCRs) convert a remarkable diversity of extracellular stimuli into precise intracellular signaling to determine cellular properties. A fascinating property of GPCRs is that ligands with subtly different chemistry can bind the same orthosteric pocket yet produce markedly different responses that remain poorly understood^1^. Ligand responses are characterized by potency, the concentration required to produce a half-maximal response, and efficacy, the capacity of a receptor-bound ligand to promote signaling^2,3^. The maximal response measured in cells reflects efficacy together with receptor abundance and the properties of the signaling system^4^. Despite extensive study, how ligand binding leads to a specific change in receptor activity remains unresolved.

Structural and biophysical studies provide a molecular framework for understanding ligand-dependent GPCR activity. High-resolution structures revealed how ligands engage the GPCR orthosteric pocket ^5–8^. Spectroscopic and single-molecule studies extended this picture, showing that ligands of different efficacy alter receptor conformational populations and the dynamics of G-protein engagement and nucleotide exchange^9–12^. Recent structures of receptors–G protein intermediates further connect efficacy to ligand-dependent progression through the activation cycle^13–17^. Together, these studies establish that ligand efficacy results from ensemble populations and transitions among them, rather than a single ligand-bound conformation^18–20^. Yet identifying a ligand-dependent contact or conformational change does not by itself establish how that feature contributes to the cellular signaling response, or whether its contribution is shared by other ligands.

Mutations can connect biophysical observations to cellular pharmacology, but are typically used to test a few residues nominated by structural or spectroscopic studies. Such experiments test specific mechanistic hypotheses without systematically comparing functional contributions across a receptor. Therefore, for activation which likely depends on many residues, it remains difficult to establish whether partial and full agonists depend on the same activation mechanisms, which regions in the receptor are most consequential for signaling, and how these dependencies change with ligand concentration. Testing mutations across the entire receptor across ligands of different efficacy and a range of concentrations would allow their functional consequences to be compared in cells. Such an experiment brings the logic of biophysical perturbation into cellular signaling, providing a route to connect structural ensemble-based models of receptor activation with the concentration–response relationships of classical pharmacology.

Deep mutational scanning (DMS) offers a method for such inquiry by linking systematic sequence perturbations to functional measurements in cells^21–26; 25,27–32^. Mutational scanning in ligand binding proteins has revealed residue networks that connect ligand binding to distal active sites^23,33–35^. Prior GPCR mutational scanning studies have identified residues involved in ligand recognition, receptor activation, and G protein coupling^32,36–38^. However, identifying residues that support signaling under a particular condition does not establish how their contributions change across ligands of different efficacy. Interpreting these differences also requires separating several effects that can produce similar changes in a cellular response. A single ligand concentration can conflate changes in potency and maximal signaling, while reduced receptor abundance can resemble a defect in activation. Moreover, a full agonist can retain a maximal response despite a partial impairment of receptor function. Combining receptor-wide mutagenesis with measurements of concentration dependence, surface expression, and responses to ligands spanning antagonism to full agonism therefore provides a way to determine how the functional requirements for signaling change across a receptor’s pharmacological range.

Here, we use the µ-opioid receptor (µOR) as a model GPCR to determine how receptor sequence shapes ligand efficacy and potency (**Figure 1A**). Decades of discovery have identified diverse µOR ligands, including morphinans, fentanyls, peptides, and other synthetic scaffolds, that all bind the same pocket but produce distinct signaling efficacies^39,40^. This chemical and functional diversity, together with extensive structural and biophysical studies, makes the µOR well suited to examining how receptor sequence shapes ligand efficacy and potency. We developed a deep mutational pharmacology approach, measuring the effect of ∼10,000 µOR variants covering every substitution at all 399 positions in the receptor on signaling in cells against 15 opioids that span a full efficacy range. For three agonists, we obtained full concentration-response series, enabling calculation of how each variant influences efficacy and potency. These ∼400,000 variant-ligand measurements resolve where in the receptor efficacy is set, whether ligands of different efficacy depend on the same or distinct positions, and, using double mutants, how those residues are coupled. The resulting functional maps support a model in which ligands of different efficacy activate the receptor thorugh distinct mechanisms.

**Figure 1:**
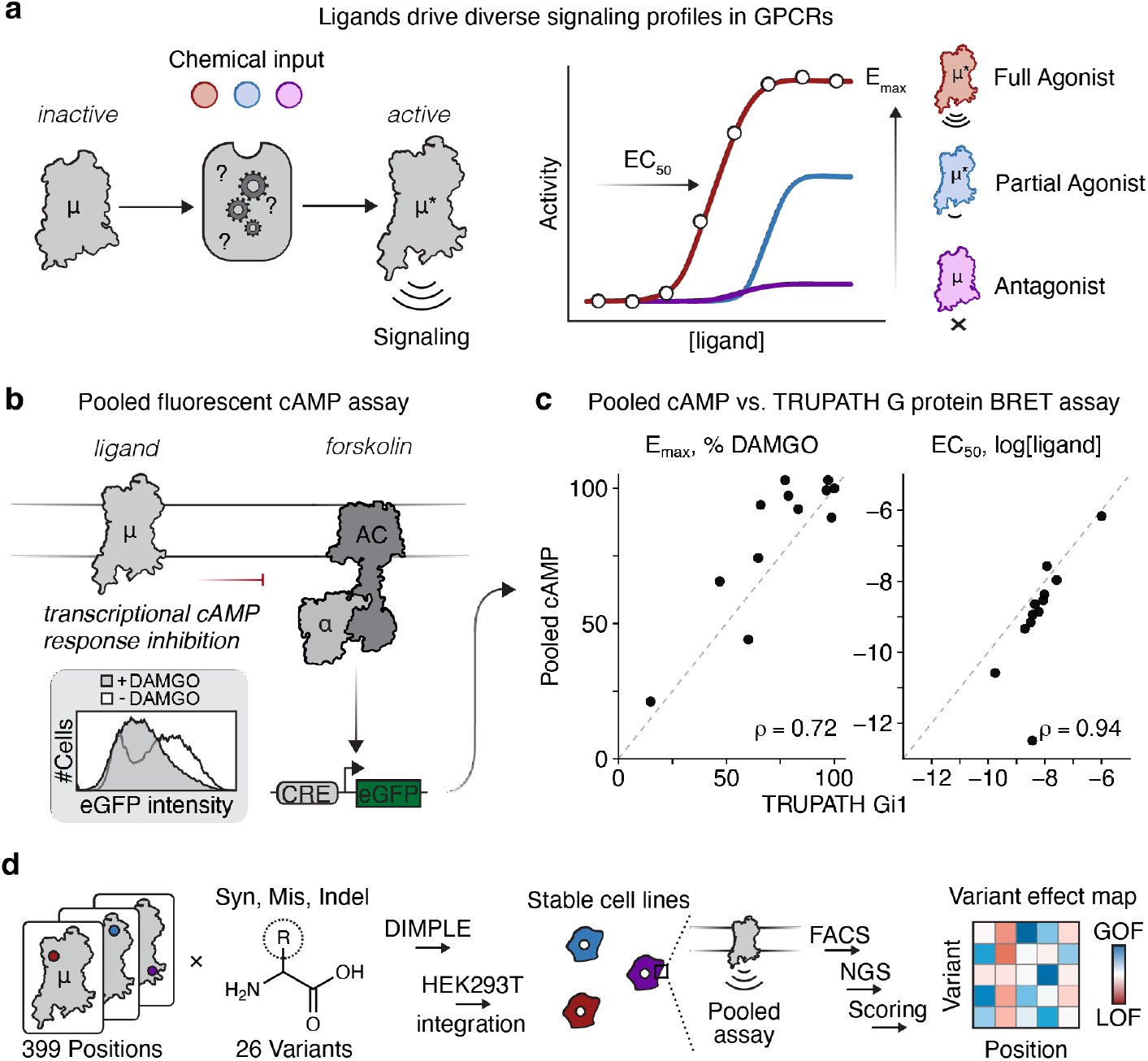
Deep mutational pharmacology. (a) GPCR activity is modulated by diverse chemical inputs. Ligands activate receptors with different efficacies and potencies. (b) Schematic of pooled cAMP inhibition assay enabling screening of large mutational libraries. Downstream cAMP is monitored by eGPF production of a cAMP response element after co-incubation with forskolin and a given receptor ligand. Example flow cytometry data shown for DAMGO (agonist) activating a variant library of µOR. (c) Comparison of efficacy (E_max_) and potency (EC_50_) parameters between pooled cAMP inhibition assay and TRUPATH Gi1 BRET for a panel of opioid ligands. Spearman coefficient is shown. The grey dashed line indicates parity between two assays. **See Supplemental Figure 1** for curves and fit parameters. (d) Abbreviated workflow of µOR mutational library generation, cell line integration, screening, and processing steps to arrive at maps of variant effects for a given assay and condition.

## Results

### Deep mutational pharmacology of µOR

The µOR couples to inhibitory G proteins (Gi, Go, and Gz) that suppress adenylyl cyclase activity^41,42^. To read out variant-level inhibitory signaling in a pooled format, we adapted a fluorescent transcriptional reporter of cAMP previously developed for mutational scanning of Gs-coupled receptors (**Figure 1B**)^32^. Here, cells are stimulated with forskolin to elevate cAMP production, which drives expression of eGFP; activation of µOR inhibits cAMP production and is detected as a reduction in reporter fluorescence (**Figure 1B**). Against a panel of reference ligands, this transcriptional assay reproduces the expected pharmacology of full agonists, partial agonists, and antagonists as evaluated by TRUPATH^43^, a well-established assay that directly measures GPCR-mediated G protein activation. For E_max_ the assays are well-correlated, and for EC_50_ the assays are nearly identical (E_max_ Spearman ρ = 0.72, EC_50_ Spearman ρ = 0.94; **Figure 1C, Supplemental Figure 1**)^44^. The cAMP reporter assay therefore captures the same pharmacological information as classic GPCR assays, while providing the throughput required to assess thousands of receptor variants.

We constructed a variant library of human µOR at all 399 positions using the DIMPLE pipeline (**Figure 1D, Supplemental Figure 2A**)^45^. Each position was covered by all possible missense substitutions, a synonymous variant, and insertions and deletions of 1, 2, and 3 amino acids (**Figure 1D, Supplemental Figure 2B**) ^46^. The library was integrated as single-copy genomic insertions in HEK293T cells via a Bxb1 landing-pad system, ensuring one variant was incorporated per cell ^47^. We observed 97.7% of designed missense variants in the final library (**Supplemental Figure 2C**).

### Potency and efficacy depend on distinct receptor regions

To resolve variant effects on ligand efficacy and potency, we assayed the µOR variant library against three agonists spanning major opioid chemotypes: morphine (alkaloid natural product), DAMGO (endogenous enkephalin peptide analog), and fentanyl (synthetic drug) (**Figure 2A, Supplemental Figure 2D-E**). All three bind the canonical orthosteric ligand pocket in µOR ^48^. We profiled the library across a range of concentrations for each ligand, scoring the activity of every variant at each concentration. We additionally profiled the library in the absence of ligand to examine how variants affect basal signaling (**Figure 2C, Supplemental Figure 2F**).

**Figure 2:**
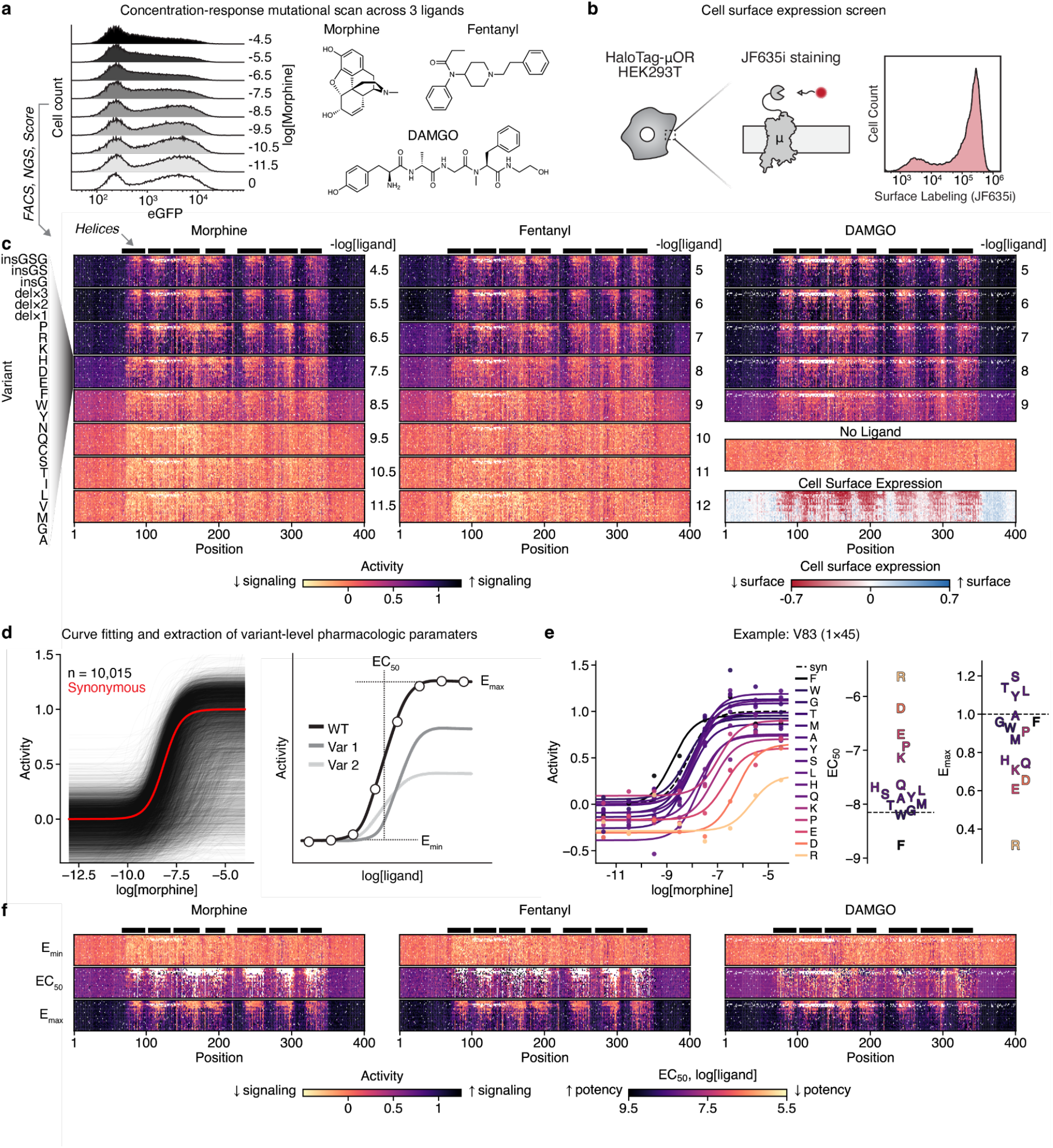
Concentration-response deep mutational scan of the µOR. (a) Example flow cytometry data of µOR mutational library activation by morphine using the pooled cAMP inhibition assay. Chemical structures of each ligand screened in concentration-response mutational scanning experiments. (b) Schematic workflow for measuring cell surface expression via staining N-terminal HaloTag with cell-impermeable dye JF-635i. (c) Stacked variant score heatmaps for concentration-response mutational scans of morphine, fentanyl, and DAMGO. Additional heatmaps for signaling in the absence of ligand and cell surface expression. Receptor position is along the x-axis, and the variant order along the y-axis is shown. Ligand concentration in M is indicated to the right of each applicable heatmap. Data and color scales for signaling assays are normalized such that the synonymous E_max_ is set to 1 and EC_50_ set to 0 for each ligand. White tiles indicate missing data. Transmembrane helices are indicated by the black bars above each heatmap section. (d) Concentration-response curve fits shown for each variant in the library in response to morphine (black). Average synonymous curve shown overlaid (red). Cartoon indicating the parameters extracted from each fit: E_min_, EC_50_ (potency), and E_max_ (efficacy). (e) Example data for position 83 variants from the morphine dataset. Individual curve fits and raw data are shown for each variant. Distributions of derived EC_50_ and E_max_ parameters for each variant are shown to the right. (f) Stacked variant heatmaps for E_min_, EC_50_, and E_max_ parameters derived from the concentration-response mutational scans for each ligand. Legends are shown at the bottom of the panel. White tiles indicate missing data or parameters that are unable to be determined for a given variant. Transmembrane helices are indicated by the black bars above each heatmap section as in **c**.

Several features of the resulting activity profiles provide confidence in the approach. At low ligand concentrations, where the receptor is expected to be minimally active, the mutational landscape is nearly flat (**Figure 2C**). Receptor activity increases with agonist concentration, and each ligand titrates at an EC_50_ concordant with its potency against wild-type µOR. Regions of the receptor known to be intolerant to mutation, such as the transmembrane helices, show loss of function regardless of agonist concentration, while more tolerant regions such as the termini and loops show minimal signaling defects. Finally, many signaling defects track with reduced cell surface expression, which was measured in a parallel screen by labeling of an N-terminal HaloTag (**Figure 2B-C**). Comparing variant scores at the highest concentration of each ligand against surface expression yields a high position-averaged correlation (Spearman ρ = 0.77-0.81 for each ligand; **Supplemental Figure 3**).

For each ligand, we fit the concentration response profiles at all ∼10,000 µOR variants to a three-parameter Hill equation, extracting each variant’s basal activity (E_min_), potency (EC_50_), and efficacy (E_max_) (**Figures 2D-F, Supplemental Figure 4**). A variant’s E_max_ reflects both how many receptors reach the cell surface and how well each receptor couples to G protein, so a reduced E_max_ can come from lower expression or from weaker coupling. To separate these, we fit an operational model^4^, widely used by pharmacologists, that uses each variant’s surface expression to resolve its per-receptor coupling (**Supplemental Figure 5**). E_max_ and per-receptor coupling remain largely correlated across the library. However, the operational model fit resolves the two contributions to E_max_, separating variants where low signal reflects low expression, with coupling near wildtype, from those with genuine coupling defects (**Supplemental Figure 5**).

Most variants reduce both efficacy and potency, but a subset selectively alters one parameter (**Figure 3A, Supplemental Figure 6**). We therefore asked which regions of µOR are most sensitive in each. We use morphine as the exemplar here and report the equivalent analyses for fentanyl and DAMGO in the supplement (**Supplemental Figure 6A-F**). Plotting the average missense effect on morphine E_max_ onto the µOR structure shows that efficacy determinants are distributed across the transmembrane bundle (**Figure 3B**), with the highest sensitivity at conserved class A GPCR activation motifs, including the CWxP, DRY, and Na^+^ site (**Figure 3B**)^49–51^. By contrast, the strongest potency (EC_50_) determinants localize to the orthosteric pocket (**Figure 3C**) concentrated within ∼7 Å of the ligand (**Figure 3C, Supplemental Figure 6G**). A secondary set of EC_50_-sensitive positions maps to the G protein cavity, consistent with agonist potency depending on G protein coupling as well as ligand pocket contacts. Because EC_50_ and E_max_ are coupled through receptor activation, many positions affect both (**Figure 3A**). Positions that diverge between EC_50_ and E_max_ are most concentrated in the ligand binding pocket for potency and across the activation motifs for efficacy (**Figure 3B-D, Supplemental Figure 6**). Similar to the finding from the operational model, the same trend is obtained when efficacy is normalized by expression, supporting the notion that distributed efficacy determinants are not simply due to folding deficits (**Supplemental Figure 6I-K**). Agonist efficacy and potency are therefore largely coupled, but diverge with potency depending on the ligand pocket and efficacy on a broader set of residues.

**Figure 3:**
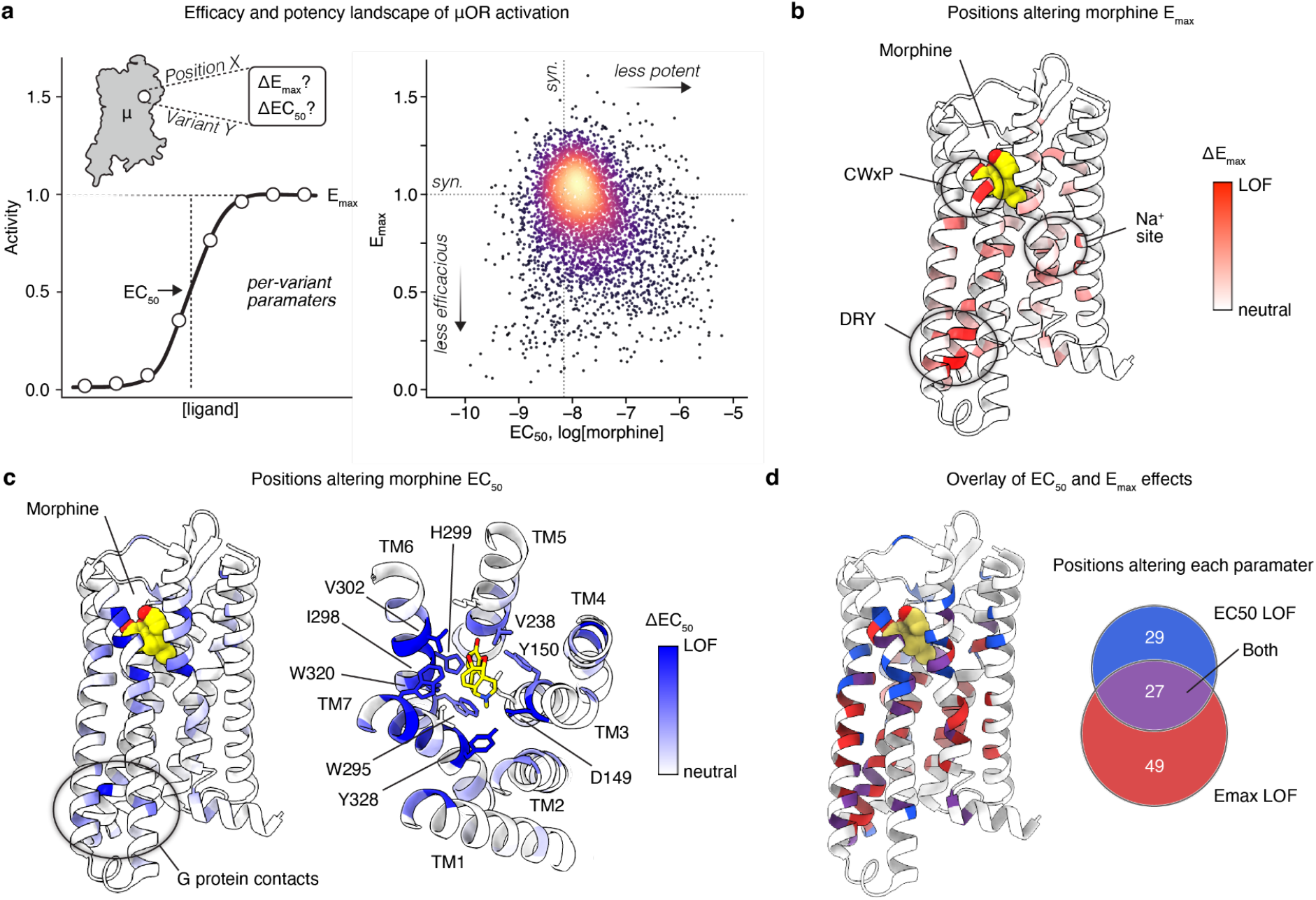
Variant effects on morphine EC_50_ and E_max_ from a DMS. (a) Landscape of per-variant EC_50_ and E_max_ effects on morphine activity. (b) Per-position average effect on E_max_ displayed on µOR-Morphine structure (PDB: 8EF6). White indicates a position behaves similar to synonymous, red indicates mutations at a given position result in decreased signaling, reflected by lower E_max_ values. (c) Per-position average effect on EC_50_ displayed on µOR-Morphine structure (PDB: 8EF6). White indicates a position behaves similar to synonymous, blue indicates mutations at a given position result in decreased signaling, reflected by lower E_max_ values. Zoom view shows interactions in the orthosteric site. (d) Overlay of per-position effects on EC_50_ (blue) and E_max_ (red) displayed on µOR-Morphine structure (PDB: 8EF6). Positions are colored if their effect on one or both parameters exceeds one standard deviation of the synonymous population.

### Agonists reweight shared potency and efficacy determinants

Are the three µOR agonists sensitive to the same mutations? We compared the mutational effects on potency and efficacy across the three ligands (**Figure 4A**). While the general mutational effects correlate between the three agonists, they are not identical (**Figure 4, Supplemental Figure 6**). Comparing morphine and fentanyl, per-position effects correlate more tightly for efficacy than for potency (Spearman ρ = 0.94 for E_max_ vs 0.63 for EC_50_, **Figure 4B, D**). To quantify how the agonists diverge, we calculated a loss-of-function bias score (LOF bias) for each pair. At each position, we first observe the effect of a given mutation on each agonist’s potency or efficacy, relative to the synonymous baseline, then compare the strongest of these effects between the two agonists. Mapping these per-position bias scores onto the receptor structure reveals where potency and efficacy determinants diverge across ligands.

**Figure 4:**
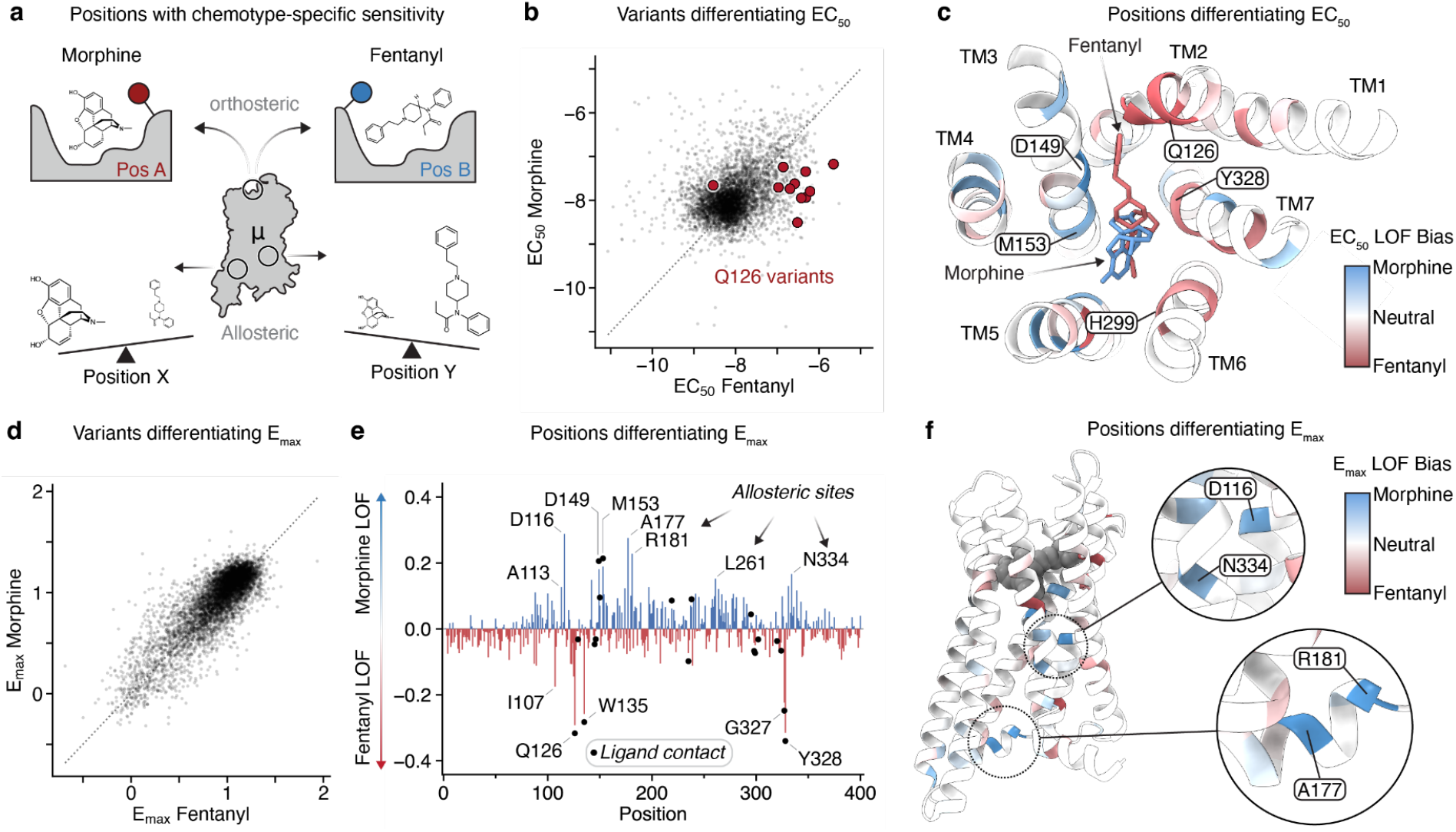
**Ligand-specific determinants of EC_50_ and E_max_** (a) Deep mutational pharmacology reveals positions with different effects on the activation of µOR by morphine and fentanyl. (b) Correlation of variant-level effects on morphine and fentanyl EC_50_ (Spearman ρ = 0.34). (c) Structure mapping (PDB: 8EF5, 8EF6) of positions with divergent effects on EC_50_. Blue indicates variants at a position shift the EC_50_ of morphine more than fentanyl. Red indicates a position has a larger effect on the EC_50_ of fentanyl relative to morphine. White are positions with equivalent effects on the EC_50_ of each ligand. (d) Correlation of variant-level effects on morphine and fentanyl E_max_ (Spearman ρ = 0.76). (e) Residuals from least-squares regression of the morphine and fentanyl position-average E_max_ parameters. Positive (blue) values indicate variants at a position have a larger effect on Emax of morphine than fentanyl. Negative (red) values indicate variants at a position have a greater effect on Emax of fentanyl than morphine. Positions contacting either ligand are indicated using black dots. (f) Structure mapping (PDB: 8EF5, 8EF6) of positions with divergent effects on E_max_. Blue indicates variants at a position shift the E_max_ of morphine more than fentanyl. Red indicates a position has a larger effect on the E_max_ of fentanyl relative to morphine. White are positions with equivalent effects on the E_max_ of each ligand. Ligands are shown in grey.

Morphine and fentanyl make distinct interactions in the µOR orthosteric pocket; we therefore expected that mutations would have different effects on the EC_50_. Indeed, positions that selectively contact one ligand show high EC_50_ LOF Bias scores. The phenylethyl moiety of fentanyl contacts Q126^2. 60^ where morphine makes minimal contact; Q126^2.60^ shows a strong fentanyl LOF Bias (superscripts denote the Ballesteros-Weinstein common numbering system for class A GPCRs^52^. Conversely, M153^3.36^ is biased towards morphine (**Figure 4B-C**). The residues that determine potency track with the distinct contacts each ligand makes in the orthosteric pocket.

The efficacy (E_max_ LOF) parameter revealed a distinct pattern (**Figure 4D-F).** Ligand specific-biases are evident in both directions, favoring morphine at some positions and fentanyl at others (**Figure 4E**). Some sites map to ligand-specific contacts, e.g. Q126^2.60^ and M153^3.36^ show E_max_ LOF biases for fentanyl and morphine, respectively (**Figure 4E**). More surprising are positions distant from the orthosteric site. Morphine is more sensitive to mutations at positions R181^34.57^ and A177^34.53^ on the intracellular loop 2 (ICL2) of the receptor and to D116^2.50^, an extensively studied residue that binds sodium ions in the inactive µOR^49,50,53–55^ (**Figure 4F**). Fentanyl is more sensitive at I107^2.41^ in TM2 (**Figure 4E**). These ligand-specific mutational dependencies remote from the orthosteric site suggest ligand-specific allosteric coupling.

### Ligand efficacy organizes mutational sensitivity across 15 ligands

Our concentration-response experiments sampled only three ligands, leaving open whether ligand-specific mutational effects recur across opioid diversity. Furthermore, the ligands we studied were all full or strong partial agonists and we wanted to understand whether mutational patterns explain differences in how diverse ligands with varying efficacy activate the receptor. We therefore conducted a mutational scan with twelve additional ligands alongside the three prior ligands at a single saturating ligand concentration (**Figure 5A-B, Supplemental Figure 1, Supplemental Figure 2G**). Single-concentration data for morphine, fentanyl, and DAMGO correlate well with the fit E_max_ parameter from the concentration-response experiments (position-mean Spearman ρ = 0.68 - 0.81, **Supplemental Figure 7A**), confirming that using data from a single saturating concentration of ligands approximates mutational effects at E_max_. The ligand panel explores two axes: 1) chemotype, spanning morphinan alkaloids, fentanyls, methadone, peptides, and synthetic scaffolds such as PZM21^56^ and SR-17018^57^; and 2) efficacy, ranging from antagonists, to partial and full agonists (**Supplemental Figure 1)**. Each ligand yields a mutational fingerprint that can be compared across the panel.

**Figure 5:**
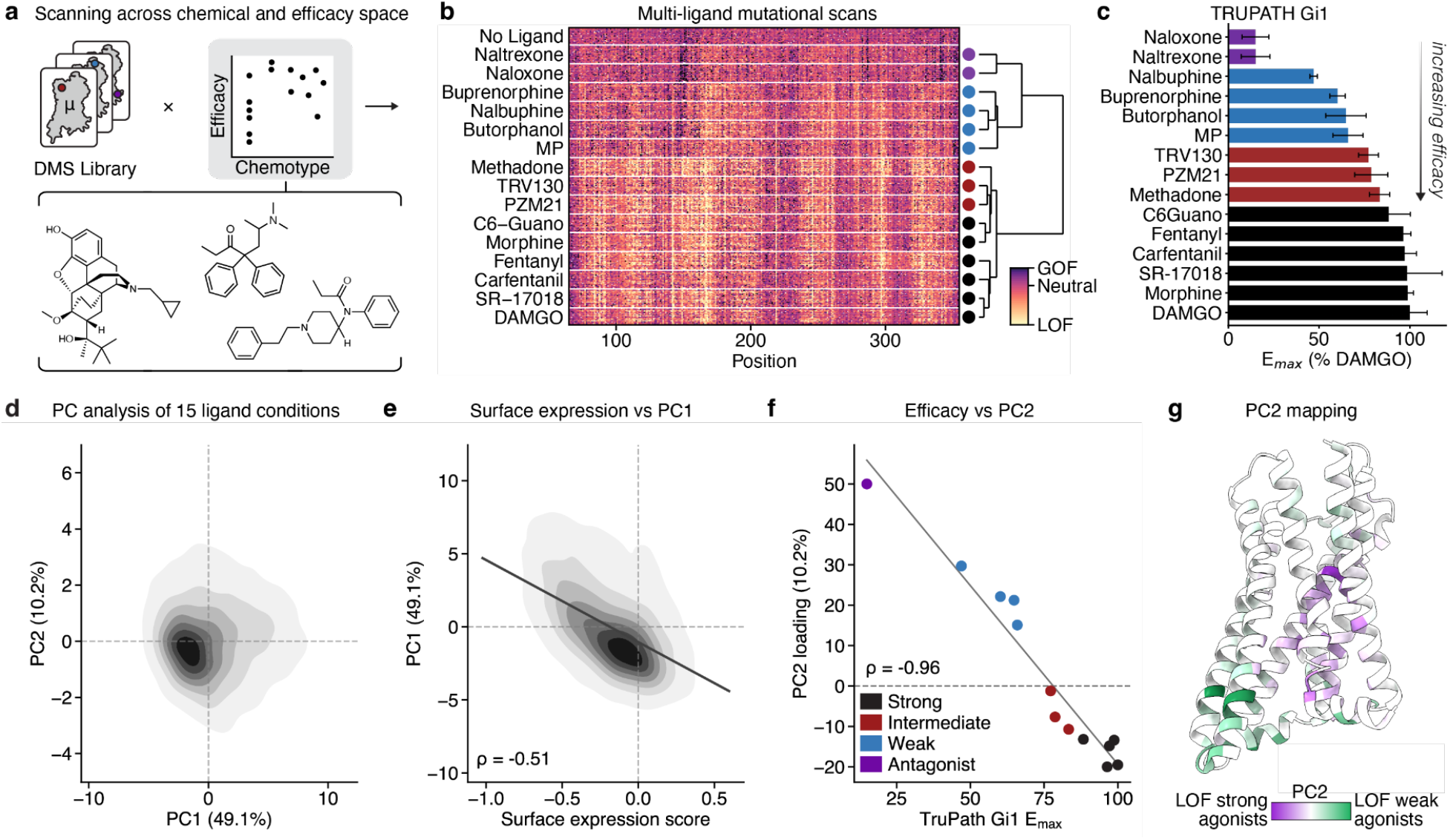
Partial agonists rely on different residues than full agonists. (a) 15 ligands were screened at saturating concentration, 10 µM. This enables comparison of mutational effects across ligands of the same chemical scaffold and different efficacy, between different chemotypes, and across efficacies. (b) Heatmap of variant effects for the untreated and 15 ligand conditions. All ligand strips are centered on the synonymous mean so that red represents decreases in signaling relative to synonymous and blue represents gain of signaling relative to synonymous. The dendrogram on the right side clusters the ligands based on average mutational effects at each position using hierarchical clustering. (c) E_max_ measurements from TruPath Gi1 assay for each ligand. Values are normalized to DAMGO (100%). Bars are colored by a ligand’s cluster membership from **b**. SR-17018 solubility was poor, so we report a single 100 µM concentration measurement here (see methods). (d) Contour plot of Principle Component (PC) 1 and 2 values across the 15-ligand dataset. (e) Contour plot of variant effects on cell surface expression compared to PC1 values. Negative surface expression scores indicate decreased expression relative to synonymous variants. Positive surface expression values indicate increased expression relative to synonymous. (f) Scatter plot of ligand PC2 loadings compared to E_max_ values from TruPath Gi1 signaling assays. (g) Structure mapping (PDB: 8EFQ) of PC2 position values. Antagonist and weak partial agonists have higher PC2 values (green). Stronger agonists have negative PC2 values (purple).

To examine how the fifteen ligands segregate by mutational scores, we clustered the per-ligand DMS effects without reference to their pharmacological labels (**Figure 5B**). Ligands cluster by relative efficacy with an order recapitulating E_max_ measured by TRUPATH (**Figure 5B-C**). We define four ligand classes: antagonists (naloxone and naltrexone), weak partial agonists (buprenorphine, nalbuphine, butorphanol, mitragynine pseudoindoxyl (MP)^58^), intermediate partial agonists (PZM21, TRV130, methadone), and strong partial and full agonists (C6-Guano^59^, morphine, fentanyl, carfentanil, SR-17018, DAMGO).

Assigning classes by TRUPATH arrives at the same groupings; that an unsupervised analysis of mutation effects recovers the efficacy ordering independently validates that efficacy is an organizing axis of the data. In the remainder of the text we use these classes as a way to probe the overall properties of ligand efficacy rather than idiosyncrasies of an individual chemotype.

What positions in µOR drive ligand efficacy? To address this question, we turned to principal component analysis (PCA), which resolves the dataset into orthogonal axes ordered by the explained variance. Two components account for ∼60% of the variance in the DMS data (**Figure 5D**). PC1 (49.1% of variance) tracks mutational effects on cell surface expression (Spearman ρ = −0.51, **Figure 5E**). Additionally, residues in the canonical class A GPCR activation motifs (e.g., DRY, CWxP, PIF, NPxxY) score highly on PC1 (**Supplemental Figure 7E**). PC1 therefore captures general activation competence rather than a source of efficacy-class or chemotype discrimination.

The ligand loadings on PC2 (which explains 10.2% of the DMS data variance) correlate with E_max_ as measured by TRUPATH (Spearman ρ = −0.96) (**Figure 5F**). PC2 therefore identifies positions that distinguish ligands of different efficacy. PCA describes each component in two complementary ways. Ligand loadings indicate how strongly each of the fifteen ligands contributes to a component, whereas each receptor variant receives a score describing its position along that component. Aggregating PC2 scores by each position and mapping them onto the structure shows that the receptor separates into two zones that display inverted mutational tolerances across ligand classes (**Figure 5G**). Positions surrounding the orthosteric site and in transmembrane helices 1-3 (TM1-3) are more sensitive to mutations for intermediate and strong agonists. Positions in the intracellular regions of TM5-6, ICL2-3, and helix 8 are more sensitive to mutations for weak partial agonists and antagonists. Sensitivity to mutation shifts across the receptor as ligand efficacy changes, suggesting that weak agonists rely on distinct regions of the receptor than stronger agonists.

### Orthosteric contacts are required in proportion to ligand efficacy

The residues a ligand contacts in the receptor binding pocket could be important for binding, signaling, or both. We use mutational data to compare how each ligand-contacting residue contributes to signaling, and whether all ligands rely on similar contacts for receptor activity.

We start with ligand-contacting residues as defined by experimental structures or computational predictions. Ten ligands have existing structures^44^. Buprenorphine, a clinically important partial agonist whose safety is attributed to submaximal μOR activation^60^, has not been resolved structurally, so we determined a 2.9 Å cryo-EM structure of the μOR–buprenorphine–miniGαi/Gβ/Gγ complex (**Figure 6A, Supplemental Figure 8**). The conformation of buprenorphine-bound μOR is similar to prior active state structures of the receptor. The binding pose of the morphinan core of buprenorphine superimposed with morphine with an RMSD of 0.648 Å (**Supplemental Figure 7B, 8**)^61^. Carfentanil also lacks an experimental structure, so we used contacts from a close analog, lofentanil, which does have an experimental structure^44^. For the remaining ligands (methadone, naltrexone, nalbuphine, and butorphanol) we predicted poses using Chai-1; for the morphinan cores, the RMSD to naloxone is 0.62-0.80 Å (**Supplemental Figure 7B-C**)^62^. With these structural models, we defined two sets of ligand-contacting residues: first-shell residues as those with any heavy atom within 4.5 Å of the ligand and second-shell residues as those within 5 Å of a first-shell residue. Overlaying these structural contacts on the ligand mutational footprints reveals which contacts are required for receptor activity.

**Figure 6:**
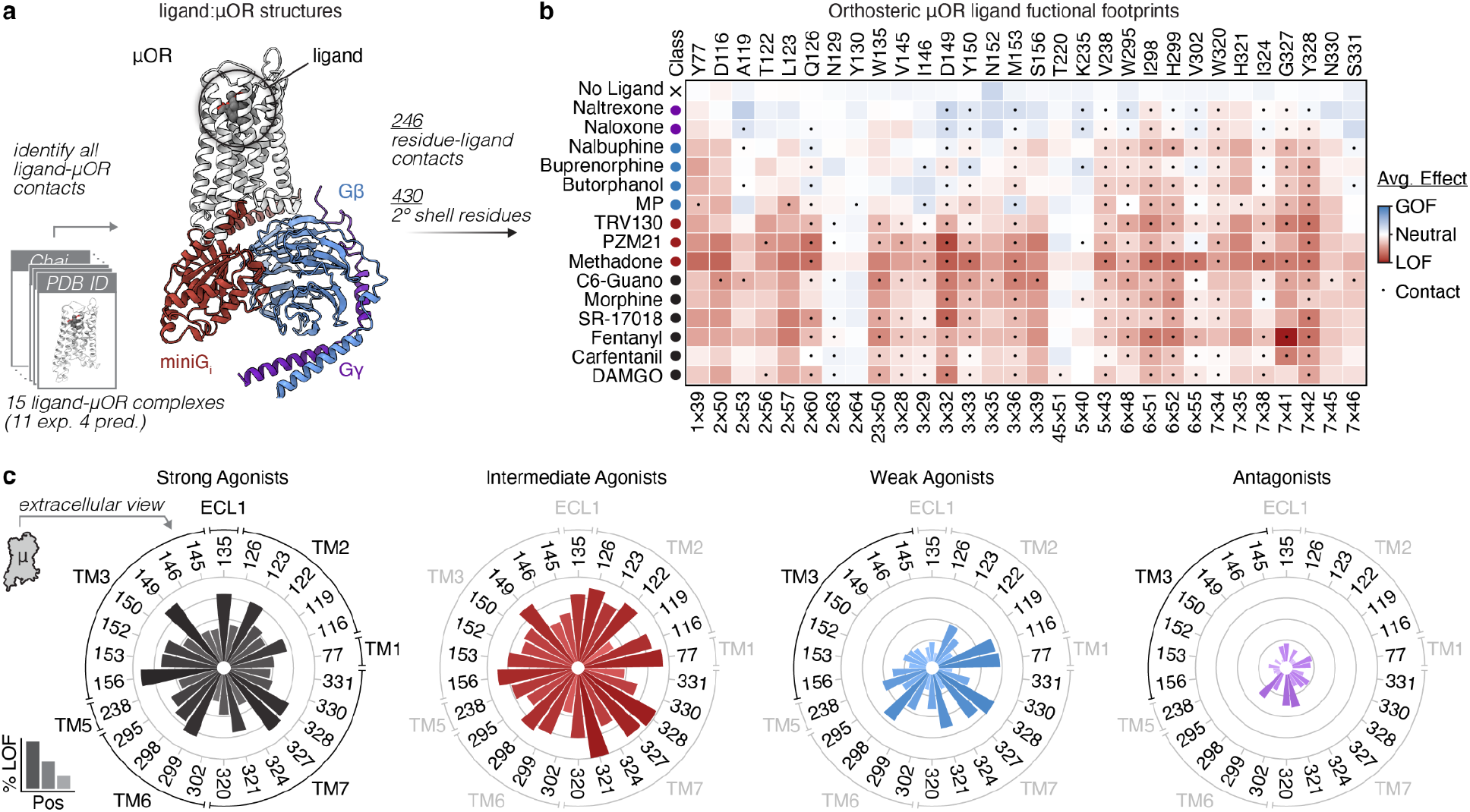
Efficacy determinants in the orthosteric pocket. (a) Workflow for identifying first and second shell residue contacts for each of the 15 assayed ligands. An atomic model is shown for buprenorphine in complexes with µOR, miniGαi, Gβ, and Gγ (PDB: 37CJ). See **Supplemental** Figure 8 for cryo-EM processing workflow and density maps. (b) Positional average missense score heatmap for all first shell contacts across the assayed ligands. Data is shown for each ligand at every position, even if there is not a direct contact. First shell contacts for each ligand are indicated by the black dots. For each ligand, blue indicates increased signaling relative to synonymous, white is centered at the synonymous mean, and red represents decreased signaling relative to synonymous. Ligand clusters determined in Figure 5B are indicated. See **Supplemental** Figure 7 for a heatmap of second shell ligand contacts. (c) Radar plots showing the average positional loss-of-function scores for orthosteric residue in each ligand cluster. Positions from **b** are shown radially, and the corresponding structural region is indicated. The bars in each plot correspond to the fraction of loss-of-function variants at a given position across the ligands in each class, defined as the fraction of variants that have scores lower than the mean synonymous minus one standard deviation of the synonymous distribution. Each ring on the plot corresponds to a 25% increment starting at 0 and ascending to 100.

Not all receptor–ligand contacts are required for activation, and the importance of some contacts depends on ligand efficacy. Weak agonists make many contacts where mutation causes minimal loss of function. By contrast, mutating those same residues causes intermediate and strong agonists to lose efficacy. This is most evident in TM3. Every opioid in our analysis contacts TM3 via D149^3.32^, a conserved position that anchors the protonated amine shared by most opioid chemotypes. TM3 is known to be critical for transmitting ligand binding to receptor activation across class A GPCRs^51^. Strong and intermediate agonists lose signaling when any TM3 position is mutated, whereas weak agonists are unaffected (**Figure 6B, Supplemental Figure 7D**). This loosened constraint suggests that weak agonists do not engage orthosteric site residues to the same extent as intermediate or strong agonists, potentially because they activate the receptor via an alternate mechanism.

### Partial agonists depend on intracellular residues that full agonists do not

Our PCA analysis also identified the intracellular site as a key differentiating region between weak agonists and intermediate/strong agonists. Intracellular residues may be important for signaling because they stabilize G protein interactions in the canonical active state^48^, in recently reported intermediate states^15,16^, or in states not resolved experimentally^63^ (**Figure 7A-B**). We collated all intracellular residues, including every position contacted by a G protein in experimental structures, and filtered for positions with functional effects in the scan. The resulting interaction map highlights intracellular residues that are important for signaling. We compare these across different ligands and with observed structural states (**Figure 7C**).

**Figure 7:**
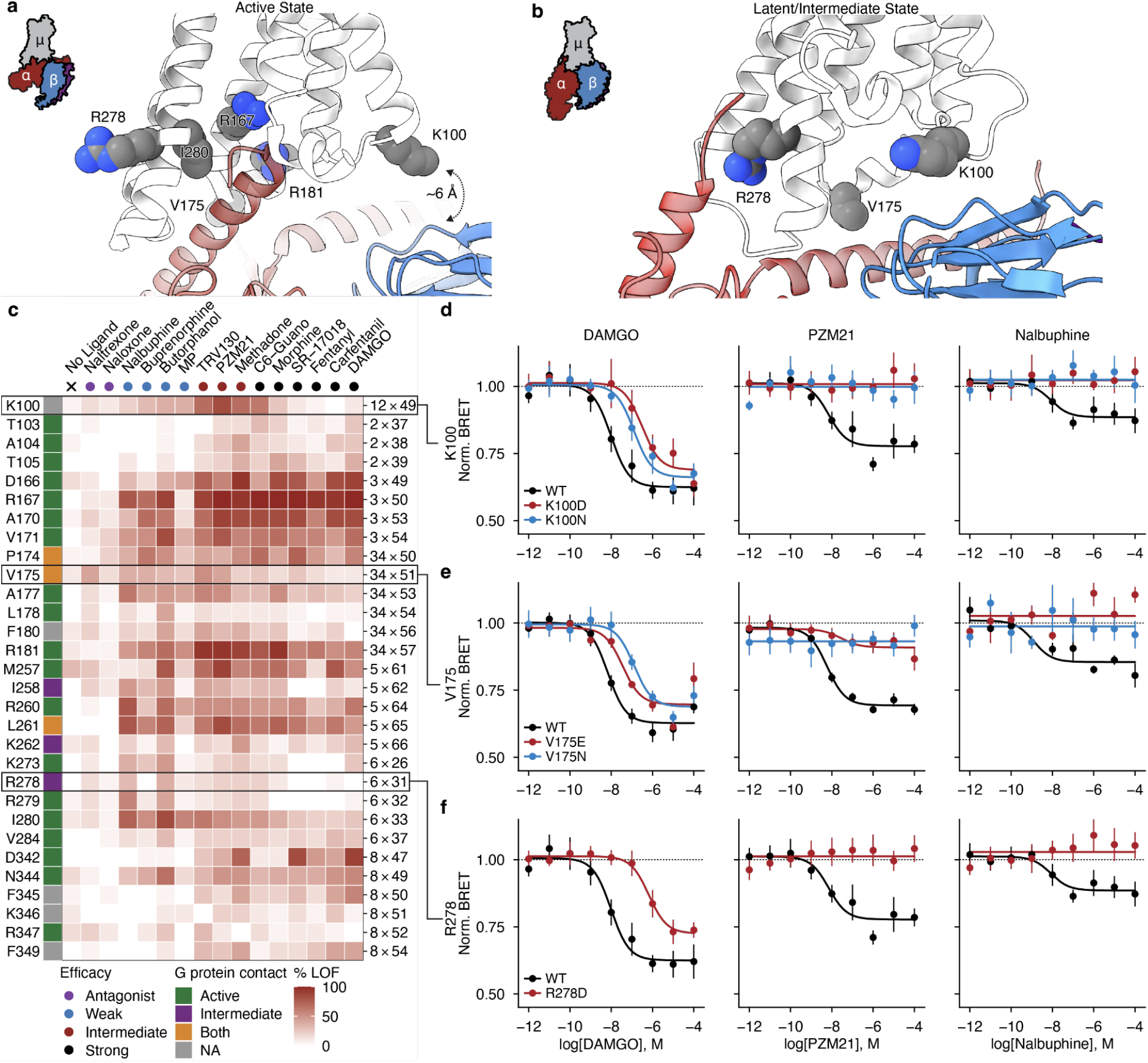
Mutations to intracellular residues have efficacy-dependent effects. (a) Zoom view of canonical active state (PDB: 8EFQ) showing µOR residues interacting with Gi heterotrimer. (b) Zoom view of latent state (PDB: 9PXW) showing µOR residues interacting with Gαi C-terminus bound extrahelically. (c) Heatmap of G protein-GPCR residue contacts along with non-contacting intracellular residues displaying the fraction of loss-of-function variants at each position for each ligand, defined as lower than two standard deviations from the synonymous mean. Data is shown for each ligand at every position. White indicates no loss-of-function variants occur for that ligand-position combination, red indicates that all variants at a position are loss-of-function. Ligand efficacy clusters determined in Figure 5B are indicated along the top axis, G protein contact class is shown across the top y-axis for each position. (d) TruPath Gi1 signaling data for DAMGO, PZM21, and nalbuphine comparing effects of aspartate and asparagine mutations at position K100^12.49^ to wildtype µOR. (e) TruPath Gi1 signaling data for DAMGO, PZM21, and nalbuphine comparing effects of glutamate and asparagine mutations at position V175^34.51^ to wildtype µOR. (f) TRUPATH Gi1 signaling data for DAMGO, PZM21, and nalbuphine comparing effects of aspartate mutation at position R278^6×31^ to wildtype µOR.

One set of residues is required regardless of ligand efficacy. The conserved DRY motif (D166^3.49^, R167^3.50^, Y168^3.51^) is loss-of-function for every ligand tested, consistent with its established role in active receptor binding to G protein (**Figure 7C**). Position R181^34.57^ in ICL2 is likewise loss-of-function across all ligands, in agreement with the human variant R181C^34.57^, which abolishes signaling (**Figure 7C**)^64^. These positions constitute a shared, efficacy-independent coupling interface.

A second set is selectively required by partial agonists (**Figure 7C**). V175^34.51^ contacts the G protein in both the active and intermediate states, and has a higher fraction of loss-of-function variants for partial agonists than for full agonists. In the TRUPATH assay, V175E/N^34.51^ reduced DAMGO potency while retaining full activation, but completely eliminated signaling by the intermediate agonist PZM21 and the weak agonist nalbuphine (**Figure 7E**). R278^6.31^, which contacts the G protein only in structural studies capturing latent/intermediate states^15,16^, behaves similarly (**Figure 7F**). K100^12.49^ in ICL1 is functionally important for partial-agonist signaling yet does not contact the G protein in any structure observed to date. Indeed, K100D^12.49^ and K100N^12.49^ shifted DAMGO potency slightly while completely disrupting partial-agonist signaling (**Figure 7D**).

A caveat for mutational studies of GPCRs is receptor reserve. Because cells often express more receptors than a maximal response requires, a full agonist can reach its ceiling even when many receptors are lost. A partial agonist, by contrast, produces less signal per receptor and therefore mutations that affect surface expression may have larger effects on efficacy. In the pooled scan, a mutation that lowers surface expression could therefore appear to selectively impair partial agonists by shrinking this reserve rather than by altering coupling. All variants tested in follow-up dose-response experiments expressed at levels comparable to the wild-type receptor by cell-surface flow cytometry (**Supplemental Figure 9**), confirming that the partial-agonist-selective differences in signaling reflect altered coupling rather than receptor abundance.

Intracellular determinants of signaling therefore depend on ligand efficacy, and partial agonists require residues that full agonists do not and potentially activate the receptor through a distinct mechanism. K100^12.49^ is the most striking: it is required for partial-agonist signaling yet makes no contact with the G protein in any resolved structure (**Figure 7A-B**).

### Functional residue networks reweight with decreasing efficacy

Full agonists are more mutationally sensitive where ligands bind while partial agonists are more sensitive where the G protein binds. To understand how the two regions are connected, we focused on the residue networks that couple binding to activation.

For each ligand efficacy class, we build a functional residue network in which nodes are positions that are loss-of-function for signaling, and edges connect nodes that have Cα atoms within 8 Å in the active receptor conformation (**Figure 8A, Supplemental Figure 10A-B**). Edges therefore mark spatial contiguity among mutation-sensitive residues, not energetic coupling. Each network reports which spatially clustered, functionally important residues a class of ligands depends on. We visualize these on the active structure of µOR.

**Figure 8:**
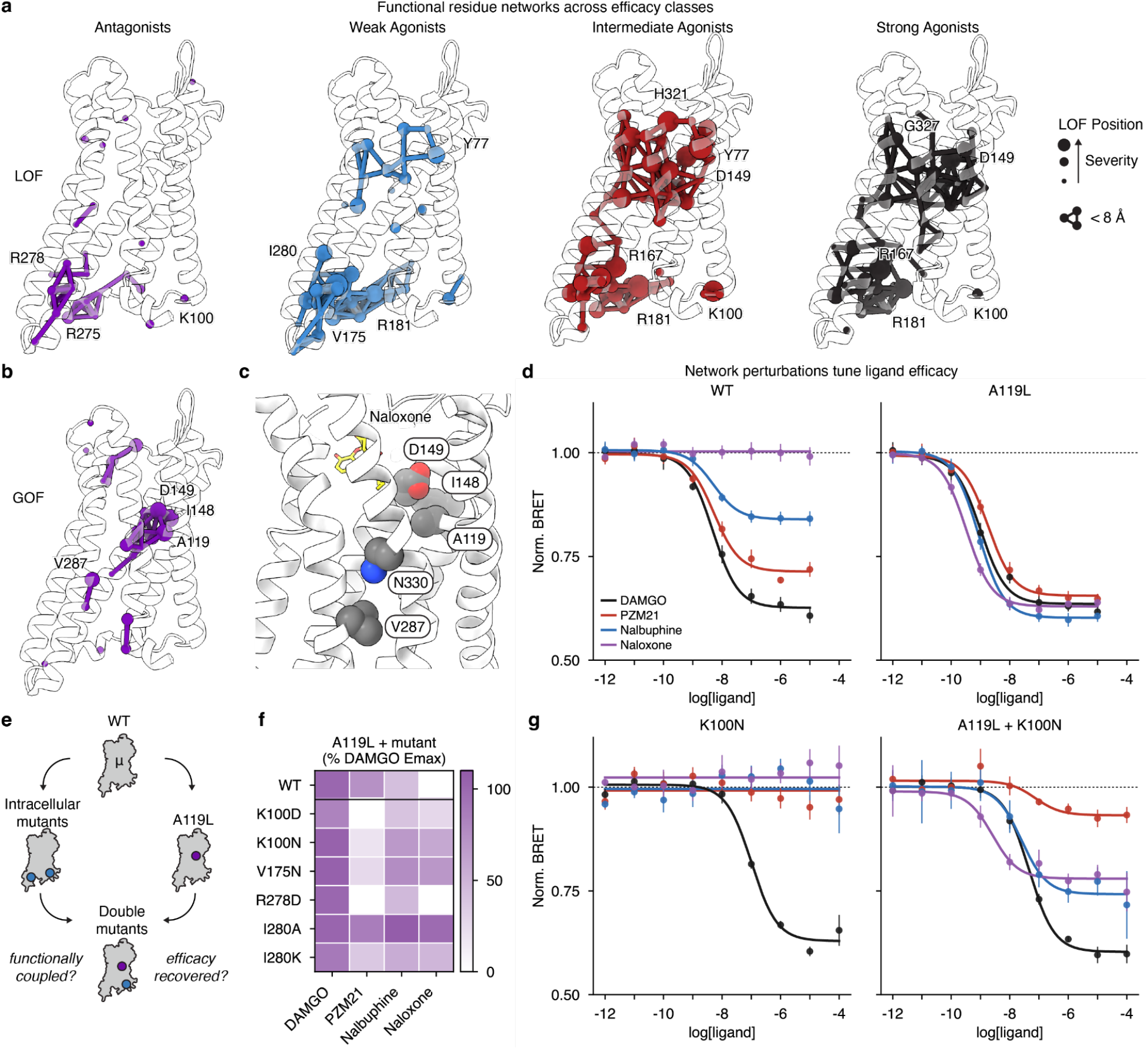
Residue networks differ across ligand efficacy. (a) Loss-of-function (LOF) residues networks across ligand efficacy. Nodes represent LOF positions and are scaled by the magnitude of LOF such that more LOF positions have larger nodes. Less severe LOF positions have smaller nodes. Connecting lines are drawn between nodes if they are 8Å or closer in 3D space (PDB: 8EFQ). (b) Gain-of-function (GOF) residues network for antagonists. Nodes represent GOF positions and are scaled by the magnitude of GOF such that more GOF positions have larger nodes. Less severe GOF positions have smaller nodes. Connecting lines are drawn between nodes if they are 8Å or closer in 3D space (PDB: 8EFQ). (c) Zoom view of residues involved in the antagonist GOF network (PDB: 9PXU). (d) TRUPATH Gi1 signaling data for DAMGO, PZM21, nalbuphine, and naloxone comparing effects of A119L^2.53^ mutation to wildtype µOR. (e) TRUPATH Gi1data assessing effect of A119L^2.53^ and K100N double mutation µOR signaling for DAMGO, PZM21, nalbuphine, and naloxone. (f) Heatmap summarizing the effect of A119L rescue of µOR signaling across several mutational backgrounds as measured by TRUPATH Gi1assay. (g) TRUPATH Gi1data assessing effect of K100N^12.49^ single mutant and K100N^12.49^-A119L^2.53^ double mutant on µOR signaling for DAMGO, PZM21, nalbuphine, and naloxone. (f) Heatmap summarizing the effect of A119L rescue of μOR signaling across several mutational backgrounds as measured by TRUPATH Gi1assay.

The networks differ across ligand efficacy in extent, location, and connectivity (**Figure 8A, Supplemental Figure 10C-D**). Strong agonists form a single, dense network spanning the orthosteric pocket, the transmembrane core, and the G protein interface. Intermediate agonists produce a network of the same shape and slightly higher density. This difference reflects sensitivity rather than the architecture of the network: strong agonists still activate the receptor despite mutations that weaken coupling, whereas intermediate agonists are more sensitive to such perturbations.

The weak agonist network is distinct from the intermediate and strong agonist networks. Density in the orthosteric pocket is decreased, consistent with weak agonists tolerating mutation of ligand pocket contacts. This is particularly clear in TM3, where strong agonists are sensitive but weak agonists are not. The nodes bridging the ligand pocket to the intracellular face are absent for the weak agonists, so what remains is an intracellular cluster rather than a continuous path. This intracellular cluster contains positions required for partial agonist signaling, including V175^34.41^, R278^6.31^, and K100^12.49^ (**Figure 7C**). Antagonists are the endpoint of this trend: their network is sparser still and confined to the G protein-proximal region.

The shape of the functional residue networks explains the observations assessing mutational effects at the orthosteric and G protein binding sites. Density of the network reflects how close a ligand operates relative to the receptor’s activation ceiling, and location reflects which part of the receptor is limiting. Strong and intermediate agonists depend on a network that spans the ligand pocket and intracellular face. As efficacy decreases, the network contracts from the pocket towards the G protein interface, and the intracellular face becomes the most sensitive. Therefore, weak agonists are most sensitive to intracellular site perturbations but resistant to orthosteric site mutations.

### A single mutation increases the efficacy of all ligands

A rigorous test of mechanism is to engineer new activities; we therefore looked for positions where mutation increases ligand-dependent signaling rather than reducing it. In the antagonist loss-of-function network, functional residues cluster on the cytosolic face, with no density and connectivity in the ligand binding site and mid-membrane region (**Figure 8, Supplemental Figure 10C-D**). The antagonist gain-of-function residue network, however, reveals a cluster of residues centered in the mid-membrane section below the orthosteric site (**Figure 8B-C**). Mutating residues in the mid-membrane region may therefore change how the receptor is activated in a way that converts antagonists into agonists.

Filtering for positions where gain of signaling is not simply elevated basal receptor activity or increased expression identified A119^2.53^ (**Supplemental Figure 7D, 9**). Structurally, A119^2.53^ bridges the ligand-binding site and the sodium site (D116^2.50^), a position where mutations are known to alter ligand activity^49,50^. Our LOF networks for intermediate and strong agonists contain A119^2.53^ as a node. Together, these observations suggest that perturbations at A119^2.53^ may modulate ligand efficacy.

Across all variants at position 119^2.53^, leucine gave the largest gain in signaling for antagonists (**Supplemental Figure 11B**). We generated the A119L^2.53^ mutation and measured signaling across the ligand panel using TRUPATH while controlling for receptor expression levels (**Supplemental Figure 9**). A119L^2.53^ converts antagonists (naloxone and naltrexone) and partial agonists (nalbuphine and PZM21) into full Gi1 agonists that match DAMGO efficacy (**Figure 8D**). Critically, A119L^2.53^ is not constitutively active, indicating that the mutation does not stabilize the active state of the unliganded receptor but instead raises the efficacy achieved once a ligand is bound. The broad reweighting of ligand efficacy with the A119L^2.53^ mutation demonstrates that the functional residue networks provide insight into the allosteric mechanisms that govern ligand efficacy.

### Mutant cycle reveals ligand-specific activation mechanisms

The preceding sections identified receptor positions where mutations alter the efficacy of strong, weak, or intermediate agonists (**Figure 7**) or raise the efficacy of all ligands (**Figure 8D**). These changes in efficacy could be explained by two models: 1) mutations change how strongly the receptor responds to a ligand, but the rank order of ligand efficacy remains the same or 2) mutations act asymmetrically on different ligands, reordering ligand efficacy. Distinguishing between these models would support whether different ligands use shared or distinct mechanisms to activate the receptor.

The single site mutations are unable to distinguish between these models. At the A119L^2.53^ mutant, all ligands have near maximal activity and it is not possible to distinguish differences in efficacy. For intracellular site mutations (K100^12.49^, V175^34.51^, R278^6.31^), weak agonists lose efficacy but intermediate and strong agonists do not; this result could also be compatible with either model. If receptor reserve or dynamic range is limiting, any mutation degrading coupling will disproportionately affect weaker agonists; this would be consistent with a model where ligand efficacy remains the same. By contrast, if weak and strong agonists use distinct mechanisms for receptor activation, these mutations should selectively affect weaker agonists; indeed this is seen for mutations in K100^12.49^, V175^34.51^, R278^6.31^. One way to distinguish these models is to boost efficacy from a distal site while maintaining these weak agonist-specific perturbations - here we use the A119L^2.53^ mutation for this purpose (**Figure 8E**). Under the first model, restoring efficacy to intracellular site mutations should restore signaling similarly for all ligands, with a preserved ligand efficacy rank order to the wild-type receptor. If, instead, mutations reweight the functional network of the receptor, it is likely that these double mutations will lead to a reranking of ligand efficacy.

We combined A119L^2.53^ with substitutions at each intracellular site position and measured Gi activation by DAMGO, PZM21, nalbuphine, and naloxone (**Figure 8E**). Receptor expression levels were matched to prevent confounds from receptor reserve (**Supplemental Figure 12**). Although DAMGO E_max_ is similar across all mutants tested, several mutations alter the EC_50_, with intracellular site mutations showing decreased potency and A119L^2.53^ increasing potency. Because receptor expression levels are kept constant, this change in DAMGO EC_50_ reflects the intrinsic capacity of the receptor network to be activated. For example, K100N^12.49^ alone eliminates PZM21 and nalbuphine responses and shifts DAMGO potency ∼21-fold; with A119L^2.53^, DAMGO potency is slightly restored and PZM21 and nalbuphine regain signaling efficacy (**Figure 8F-G**). Mutations that are most disruptive to DAMGO EC_50_ are also most disruptive to partial agonist efficacy, e.g. K100D^12.49^, V175N^34.51^, and R278D^6.31^ (**Supplemental Figure 11)**.

More surprising is the change in efficacy seen with addition of A119L^2.53^ to intracellular site mutations. For PZM21, nalbuphine, and naloxone, A119L^2.53^ increases efficacy, but in a manner that does not preserve rank order of efficacy at the wild-type receptor. At the wild-type receptor, PZM21 reaches 73% of DAMGO E_max_, while nalbuphine is 42% of DAMGO, and naloxone is silent (**Figure 8F-G, Supplemental Figure 12**). In the A119L^2.53^ / K100N^12.49^ background, this order inverts, with nalbuphine and naloxone reaching ∼60%, while PZM21 is ∼18%. Similar reversal trends are observed when A119L^2.53^ is combined with the other intracellular site mutations, indicating that coupled mutations to the receptor network lead to non-linear transformation of ligand efficacy (**Figure 8 F-G, Supplemental Figure 12**).

That there is a reordering in ligand efficacy implies that ligands of different efficacy activate the receptor through distinct mechanisms, not by simply engaging the same allosteric pathway with different strengths. Ligand efficacy is therefore a property of the ligand and the receptor sequence together, and mutations can fundamentally retune ligand efficacy.

## Discussion

The deep mutational pharmacology results presented here demonstrate that ligand efficacy is governed by residues distributed across the receptor and that ligands activate the receptor through distinct mechanisms. Across the fifteen opioids profiled here, mutational sensitivity varies systematically with efficacy. Stronger agonists are more sensitive to perturbations in the ligand binding pocket, whereas lower efficacy agonists are more sensitive to intracellular site residues. Although potency and efficacy share many determinants, the strongest effects on potency are concentrated near the ligand-binding site, while effects on efficacy extend throughout the receptor. The distinct mutational dependencies for weak vs. strong agonists are not simply consequences of different maximal responses. A single substitution in the core of the receptor, A119L^2.53^, raises the efficacy of every ligand and converts antagonists to agonists. Combining A119L^2.53^ with distal intracellular mutations reorders the relative efficacy of different ligands.

We reach several conclusions from these observations. First, efficacy is not reducible to the chemistry of an agonist and its contacts in the ligand binding pocket; instead efficacy is determined by both the ligand and the entire receptor sequence. Small perturbations to either can have dramatic consequences for efficacy. Second, our results highlight that weak and strong agonists have distinct mutational dependencies, underscoring that agonists likely activate receptors by distinct mechanisms. Third, functional networks derived from mutational data can reveal connections that are related to the underlying allosteric pathways that govern receptor function. Our ability to identify a specific residue in which a mutation uniformly raises ligand efficacy, but does not drive constitutive activity, highlights the importance of a network-driven view of receptor function. Finally, our results with double mutations demonstrate that efficacy is not simply the result of ligands stabilizing a single continuum along two conformational states. Instead, weak and strong agonists engage the receptor with distinct dependencies suggestive of distinct allosteric mechanisms and perhaps conformational states.

The functional dependencies of efficacy revealed here extends the conformational ensemble view of efficacy developed through biophysical and structural studies of GPCRs. Spectroscopic and single-molecule studies have shown that full and partial agonists alter receptor conformational populations and the dynamics of G protein engagement and nucleotide exchange^9,10,12,19,20,65^. Differences associated with low-efficacy ligands, however, remain difficult to distinguish along a single conformational coordinate; prior studies for µOR, for example, find little change in intracellular conformations in the absence of G protein^9^. With G proteins, both structural and spectroscopic studies find evidence of distinction between weak and strong agonists^9,12,20,66^. Recent structures have captured ligand-dependent intermediates during nucleotide release and G protein activation, providing a fuller picture of receptor-G protein ensembles that underlie efficacy^15,16^. Our measurements connect this molecular picture to cellular signaling by identifying perturbations that distinguish ligand responses without first selecting a particular structural snapshot to examine. Some efficacy-sensitive positions, such as R278^6.31^, participate in interfaces observed in intermediate receptor-G protein complexes. Others, including K100^12.49^, have no direct G protein contact in the structures examined to date, although earlier NMR studies implicated ICL1 in initial stages of G protein coupling^67,68^. These positions provide specific tests of how interactions within and between receptor regions influence productive signaling. The network and ensemble descriptions are therefore complementary: the biophysical studies identify molecular states and transitions, while the mutational measurements establish where perturbations have functional consequences during a cellular signaling event. We anticipate that the functional dependencies we identify here will therefore serve to both generate new hypotheses on structural mechanisms and to test the relevance of biophysical observations.

Understanding the molecular basis of efficacy is important for designing ligands that control the extent of receptor activation. For many GPCRs, some of the most successful therapeutics are partial agonists^69,70^. For µOR, the partial agonism of buprenorphine provides analgesia without the respiratory depression common to most other clinical opioids^71,72^. The success of GPCR partial agonists has motivated efforts to tune efficacy rather than simply increase affinity or potency. Structural studies have provided important examples of how to approach partial agonism^8,44,59^. For the β2-adrenergic receptor, studies of salmeterol and salbutamol connected partial agonism to differences in a hydrogen bonding network at the ligand binding side^73,74^. Our findings place such local interactions in a broader receptor context. A ligand contact does not contribute a fixed amount of efficacy independent of the rest of the protein; instead the consequences of a contact depend on an extended set of interactions. Receptor-wide functional maps could therefore complement binding poses during ligand optimization, identifying which perturbations affect a chemical series similarly and which distinguish compounds with different efficacies. Comparing these dependencies across related receptors could likewise determine how sequence differences outside the immediate ligand binding pocket contribute to receptor subtype-specific pharmacology.

Uncovering ligand specific mechanisms required designing mutational experiments tailored for pharmacology. For three agonists, concentration response connected sequence to classic pharmacology, while expression measurements and operational modeling helped distinguish receptor abundance from coupling. Comparing ligands spanning chemotypes and efficacies revealed functional dependencies that apply to efficacy and not only to ligand chemistry. Assessing loss and gain of function then identified single and double mutations to test a network model. While single site perturbations are informative, more extensive combinatorial perturbations, fit to quantitative models, provide a route to determining how residue networks are energetically coupled.

Molecular efficacy is one instance of a general problem in studying allostery: how does a protein determine the magnitude and character of its response to a bound molecule. Much of what we know of allosteric coupling comes from indirect measurements, such as biophysical studies of conformational states or multiple sequence alignments to identify co-evolving residue networks. The approach developed here demonstrates that this question can be addressed by designing mutational experiments around protein activity. Varying molecular inputs alongside protein sequence can reveal hidden properties that are difficult to identify from a single ligand, condition, or structural state. When these measurements are interpreted through quantitative models, mutational scanning brings the logic of biophysical perturbation into cells, connecting changes within a protein to the functional responses it produces. The rich functional dependencies arising from such inquiry reveal properties that extend the biophysical view of proteins from individual conformations and contacts to residue networks that drive allosteric behavior. Such systematic measurements enable a better understanding of protein allostery and will serve as foundational data for predictive models to design ligands with defined function, interpret how sequence variants change activity to cause disease, and engineer proteins with new useful activities.

### Limitations of the study

Several limitations qualify these conclusions. While we account for differences in surface expression to the best of our ability, it remains impossible to correct for folding deficits explicitly. It is possible that some functionally important residues simply do not fold when mutated, so positions required for folding are underrepresented among the sites we identify as efficacy determinants. The efficacy axis of the mutational scanning data is set by the readout from a transcriptional cAMP reporter, which is amplified and lies several steps downstream of receptor-G protein coupling. As a result, the measurement integrates short and long-lived contributions to signaling, and may be compressed relative to more proximal measurements of activity with finer temporal resolution. Additionally, for the experiments performed at a single ligand concentration, we are unable to say definitively that the condition is saturating for all variants. We test candidate mutations identified from the DMS with a limited set of ligands. While these experiments are informative for the concept of efficacy, they may not hold for all ligands. The mutant cycle experiment explores the result of a single gain of function mutation, but whether other gain of function sites behave similarly remains untested. Finally, the network model drawn from the pattern of mutational sensitivities is not explicitly connected to energetic measurements of allosteric coupling. Whether these are related remains for future investigation.

## Supplemental Figures

**Supplemental Figure 1:**
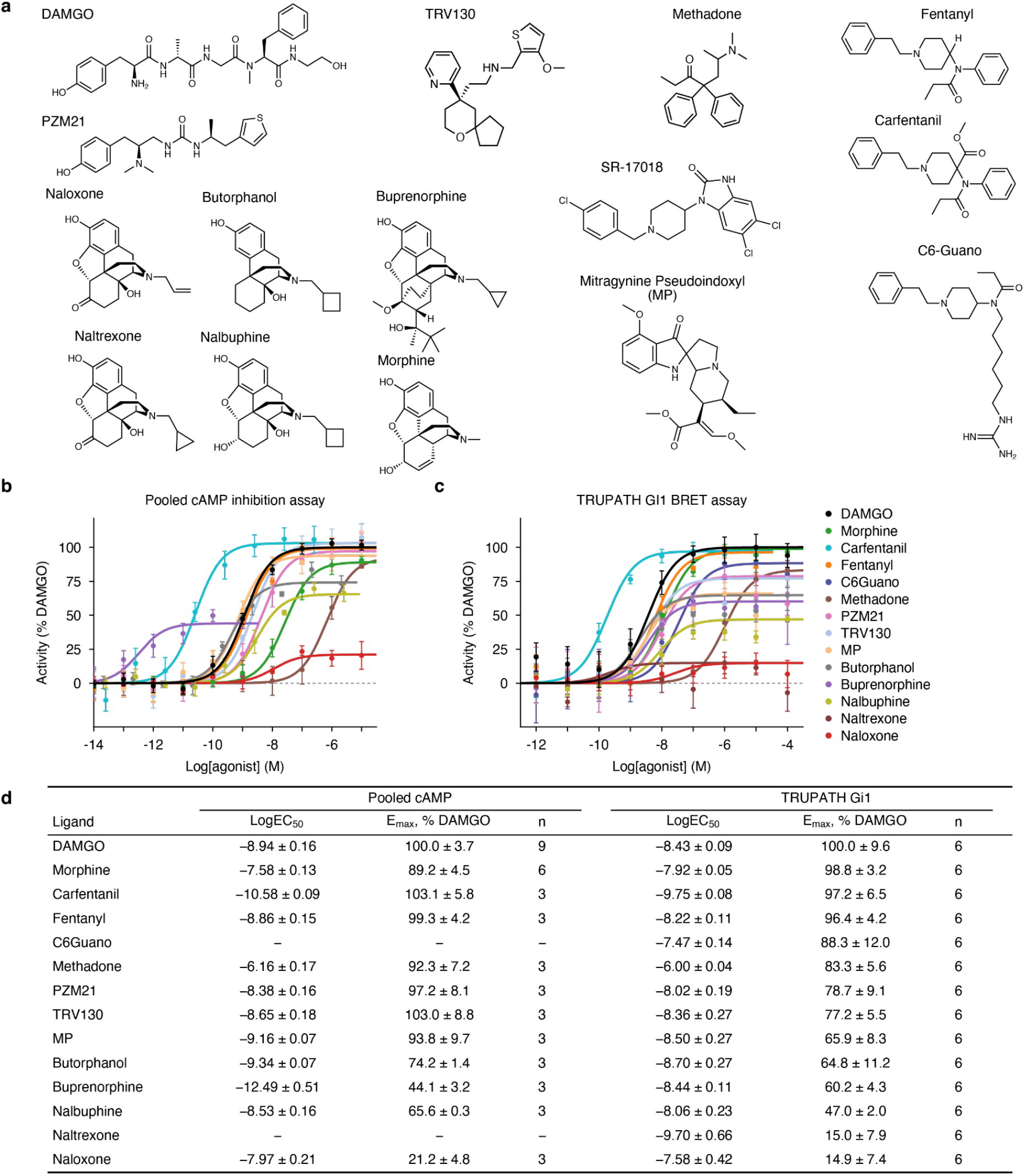
Ligands used in study and wildtype signaling data. (a) Chemical structures of all opioid ligands used in this study. (b) Concentration response curves for opioid ligands using the pooled cAMP transcriptional reporter assay on wildtype µOR. (c) Concentration response curves for opioid ligands using the TRUPATH Gi1 assay on wildtype µOR. (d) Table of pharmacologic parameters for each ligand in the pooled cAMP transcriptional reporter and TRUPATH Gi1 assays. Data shown as mean ± SEM.

**Supplemental Figure 2:**
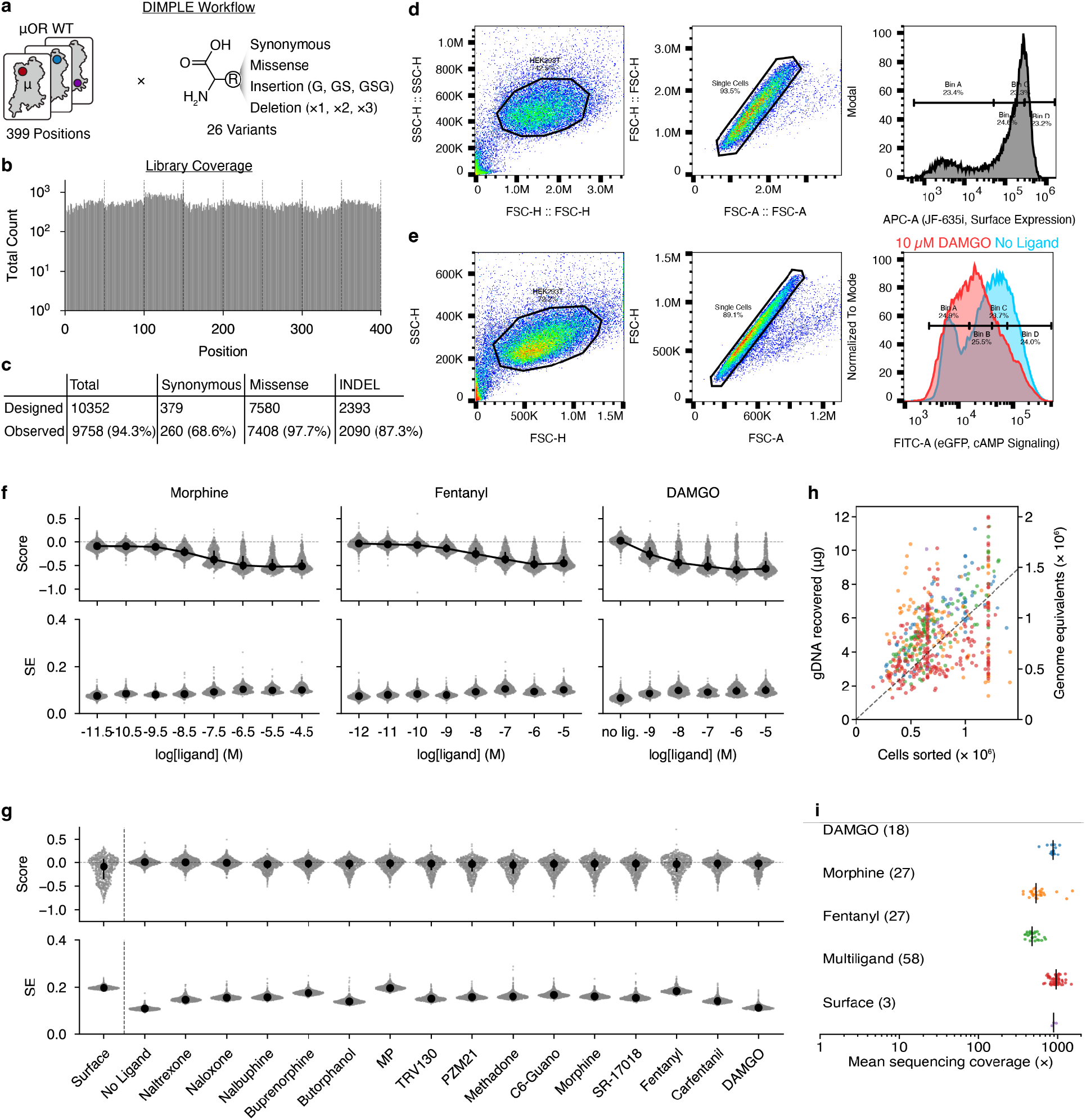
DMS library generation, screening, and QC. (a) DIMPLE workflow schematic showing variant types designed at each position of the µOR. (b) Positional coverage of the mutational library sequenced at ∼20x coverage. (c) Table of designed and observed variants in the library sequenced at ∼20x coverage. (d) Example FACS data and gating for the surface expression screens. (e) Example FACS data and gating for the signaling screens showing the library in the absence of ligand and in the presence of full agonist DAMGO at 10 µM. (f) Score and SE plots for concentration response screens with Morphine, Fentanyl, and DAMGO. Negative scores represent inhibition of cAMP signaling. Positive scores represent increased cAMP signaling. Black dot and bar represent the median and IQR. (g) Score and SE plots for the surface expression and single concentration signaling screens with multiple ligands. Synonymous variants are centered at zero. Negative values represent loss of expression or signaling. Positive values represent gain of expression or signaling. Black dot and bar represent the median and IQR. (h) Scatter plot collating the number of sorted cells per FACS sort bin and the amount of genomic DNA recovered from each sample expressed in µg of DNA and genome equivalents (assuming 6 picograms of DNA per cell). (i) Average count per variant in each condition screened, stratified by experiment type, Blue is DAMGO concentration-response screen. Orange is the morphine concentration-response screen. Green is the fentanyl concentration-response screen. Red is the multi-ligand screen at saturating concentrations. Purple is the surface expression screen.

**Supplemental Figure 3:**
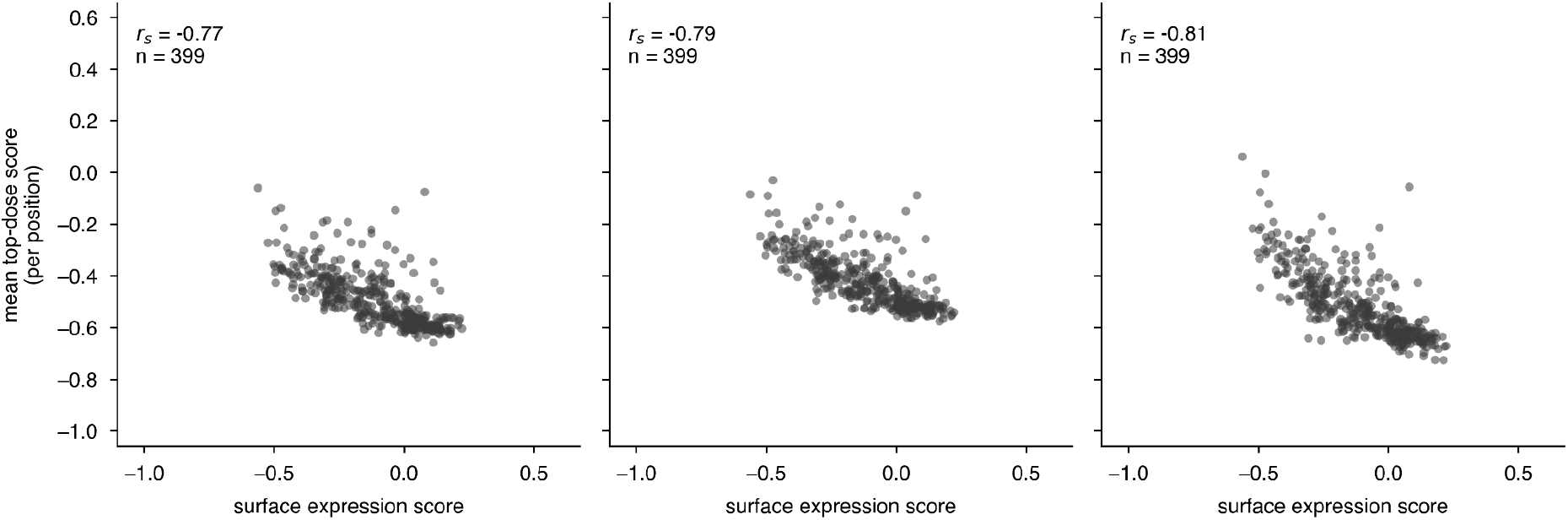
Comparison of signaling and expression effects. Position-level scatter plots of highest concentration of morphine, fentanyl, and DAMGO vs surface expression.

**Supplemental Figure 4:**
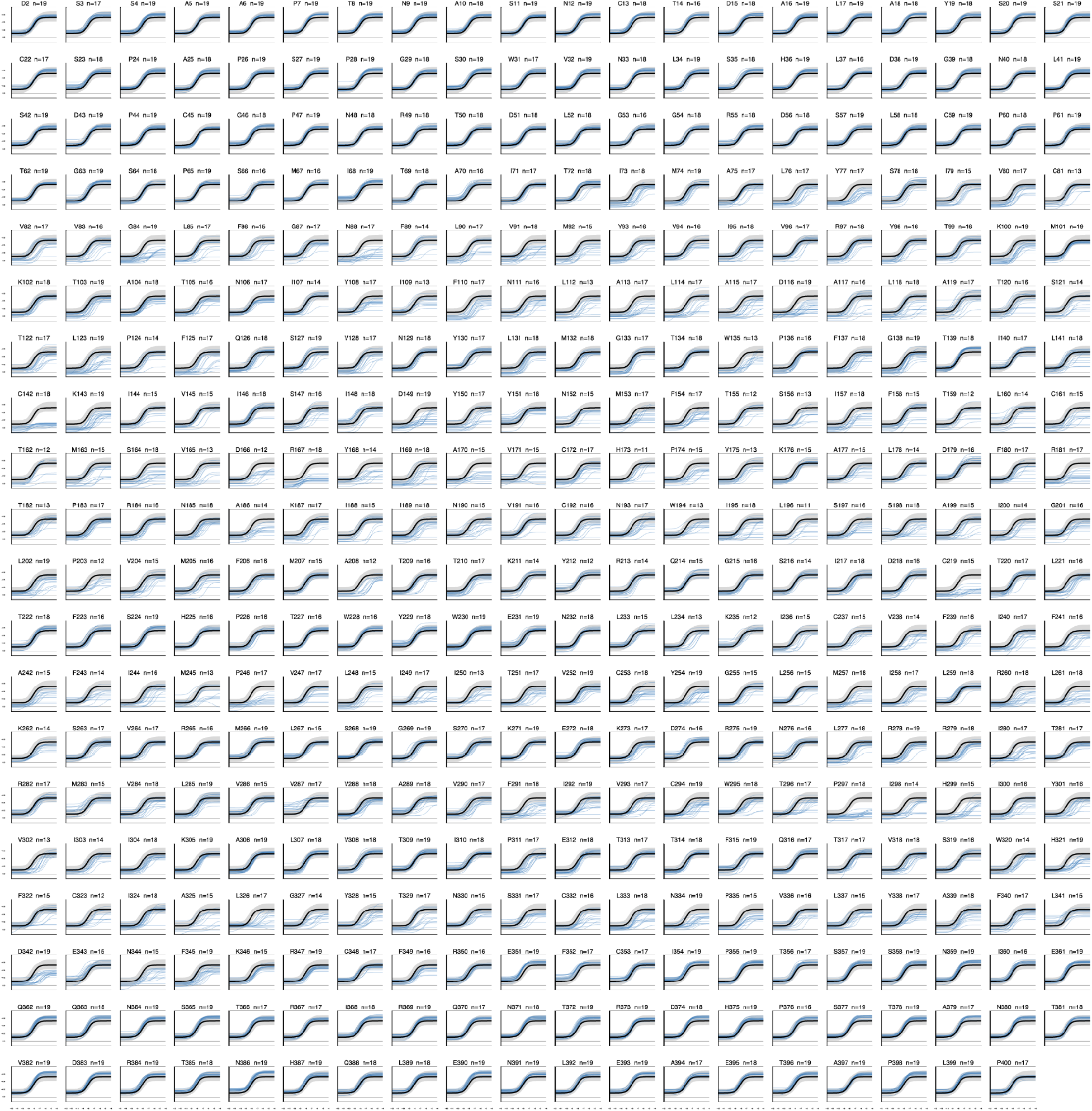
Concentration-response curves for all variant effects on morphine signaling. Per-position plots of variant concentration-response curves from the morphine screen. The black curve represents the average fit of synonymous variants in the library. The grey band is a 95% confidence interval of the synonymous variants. Blue lines are the curve fits for each variant at that position.

**Supplemental Figure 5:**
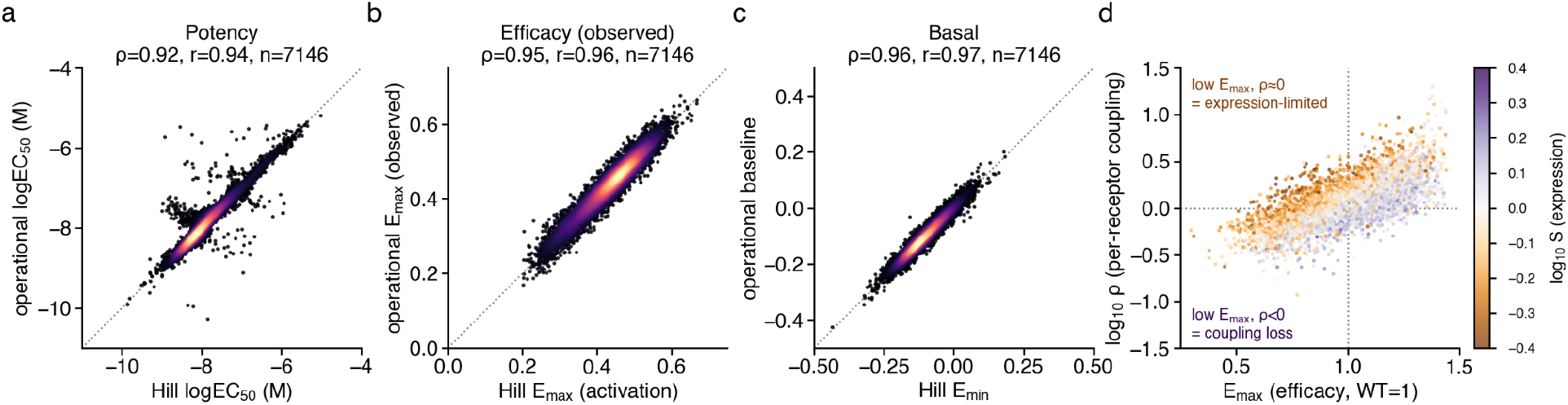
Comparison of Hill and Operational fits to concentration-response mutational scanning data. (**a-c**) Variant-level scatter plots for the morphine concentration-response mutational scan describing the relation between potency, efficacy, and baseline activity derived from Hill and Operational models. (**d**) Variant-level comparison of the E_max_ and rho parameters from the Operational model coloring by surface expression.

**Supplemental Figure 6.**
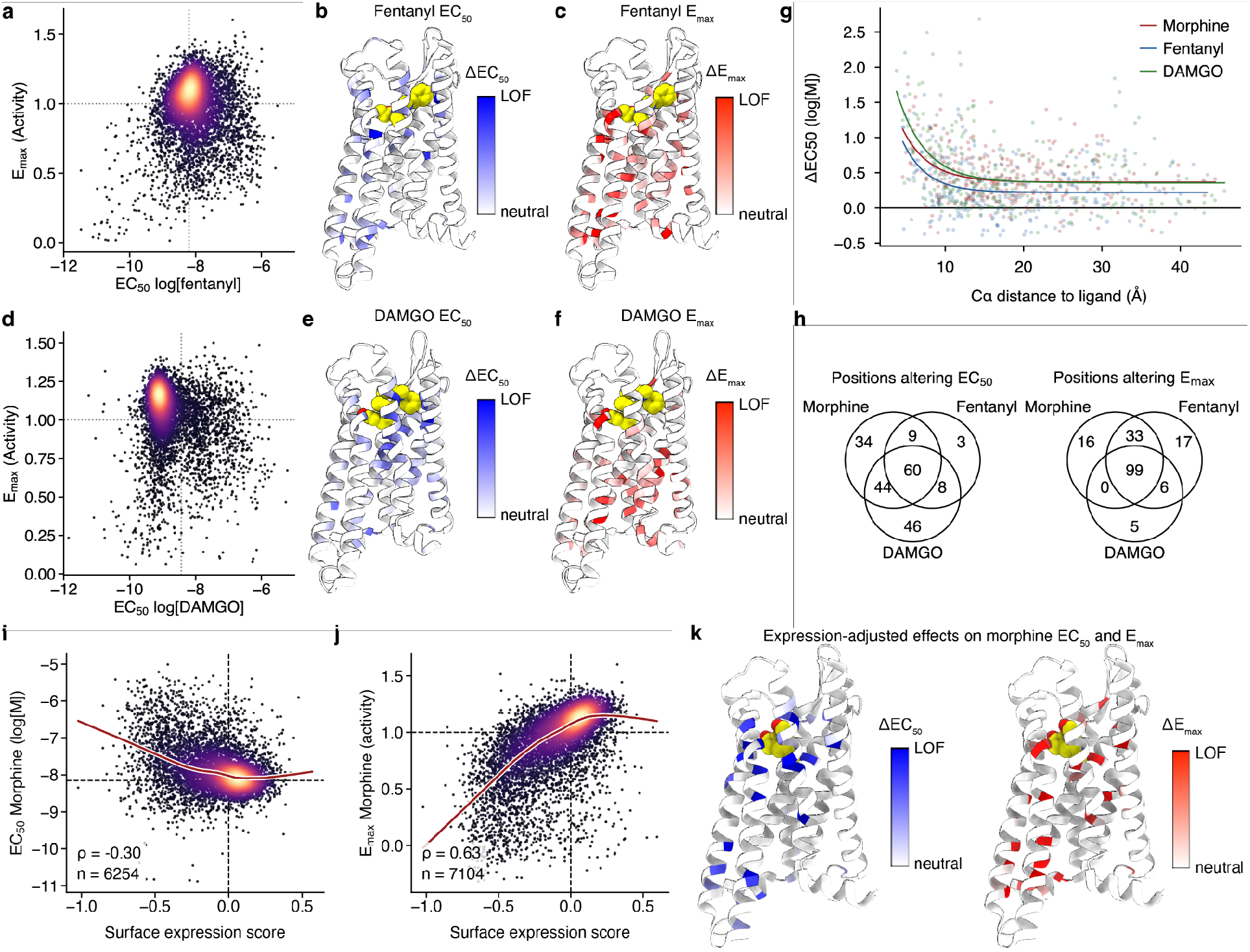
Additional EC_50_ and E_max_ data across morphine, fentanyl, and DAMGO. (a) Landscape of per-variant EC_50_ and E_max_ effects on fentanyl activity. (b) Per-position average effect on EC_50_ displayed on µOR-fentanyl structure (PDB: 8EF5). White indicates a position behaves similar to synonymous, red indicates mutations at a given position result in decreased potency, reflected by higher EC_50_values. (c) Per-position average effect on E_max_ displayed on µOR-fentanyl structure (PDB: 8EF5). White indicates a position behaves similar to synonymous, red indicates mutations at a given position result in decreased signaling, reflected by lower E_max_ values. (d) Landscape of per-variant EC_50_ and E_max_ effects on fentanyl activity. (e) Per-position average effect on EC_50_ displayed on µOR-DAMGO structure (PDB: 8EFQ). White indicates a position behaves similar to synonymous, red indicates mutations at a given position result in decreased potency, reflected by higher EC_50_values. (f) Per-position average effect on E_max_ displayed on µOR-DAMGO structure (PDB: 8EFQ). White indicates a position behaves similar to synonymous, red indicates mutations at a given position result in decreased signaling, reflected by lower E_max_ values. (g) Scatter plot of µOR residue distance from each ligand vs effects on ligand EC_50_ from the concentration-response mutational scans. (h) Venn diagrams sumarizing the number of positions which have effects on EC_50_ and E_max_ that are shared or diverge between each ligand. (i) Variant-level scatter of surface expression and morphine EC_50_. Loess fit shown in red. (j) Variant-level scatter of surface expression and morphine E_max_. Loess fit shown in red. (k) Per-position structure mapping of expression-adjusted effect on morphine EC_50_ and E_max_ displayed on µOR-morphine structure (PDB: 8EF6). White indicates a position behaves similar to synonymous, red indicates mutations at a given position result in decreased potency, reflected by higher EC_50_values.

**Supplemental Figure 7:**
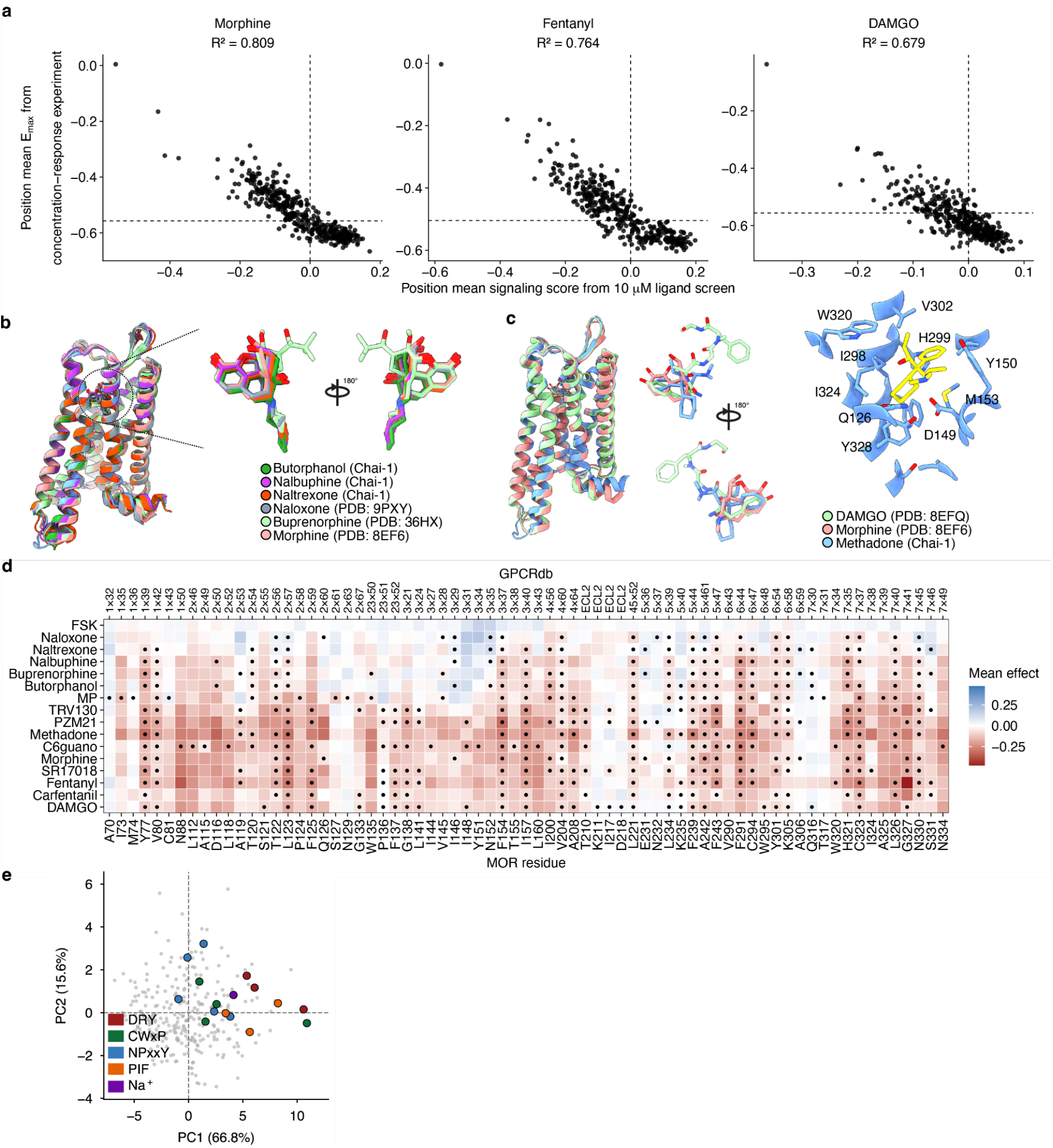
Comparison of E_max_ and saturating ligand concentration DMS scores. (a) Position average E_max_ from concentration-response experiment on morphine compared to scores from single concentration screen at 10 µM of each ligand. (b) Position average E_max_ from concentration-response experiment on fentanyl compared to scores from single concentration screen at 10 µM of each ligand. (c) Position average E_max_ from concentration-response experiment on DAMGO compared to scores from single concentration screen at 10 µM of each ligand. (d) Positional average missense score heatmap for all second shell contacts across the assayed ligands. Data is shown for each ligand at every position even if there is not a direct contact, indicated by the black dots. Blue indicates increased signaling relative to synonymous, white is centered at the synonymous mean, and red represents decreased signaling relative to synonymous for each ligand. (e) PC1 and PC2 biplot showing positions of many canonical GPCR activation motifs tend to concentrate at positive PC1 values.

**Supplemental Figure 8:**
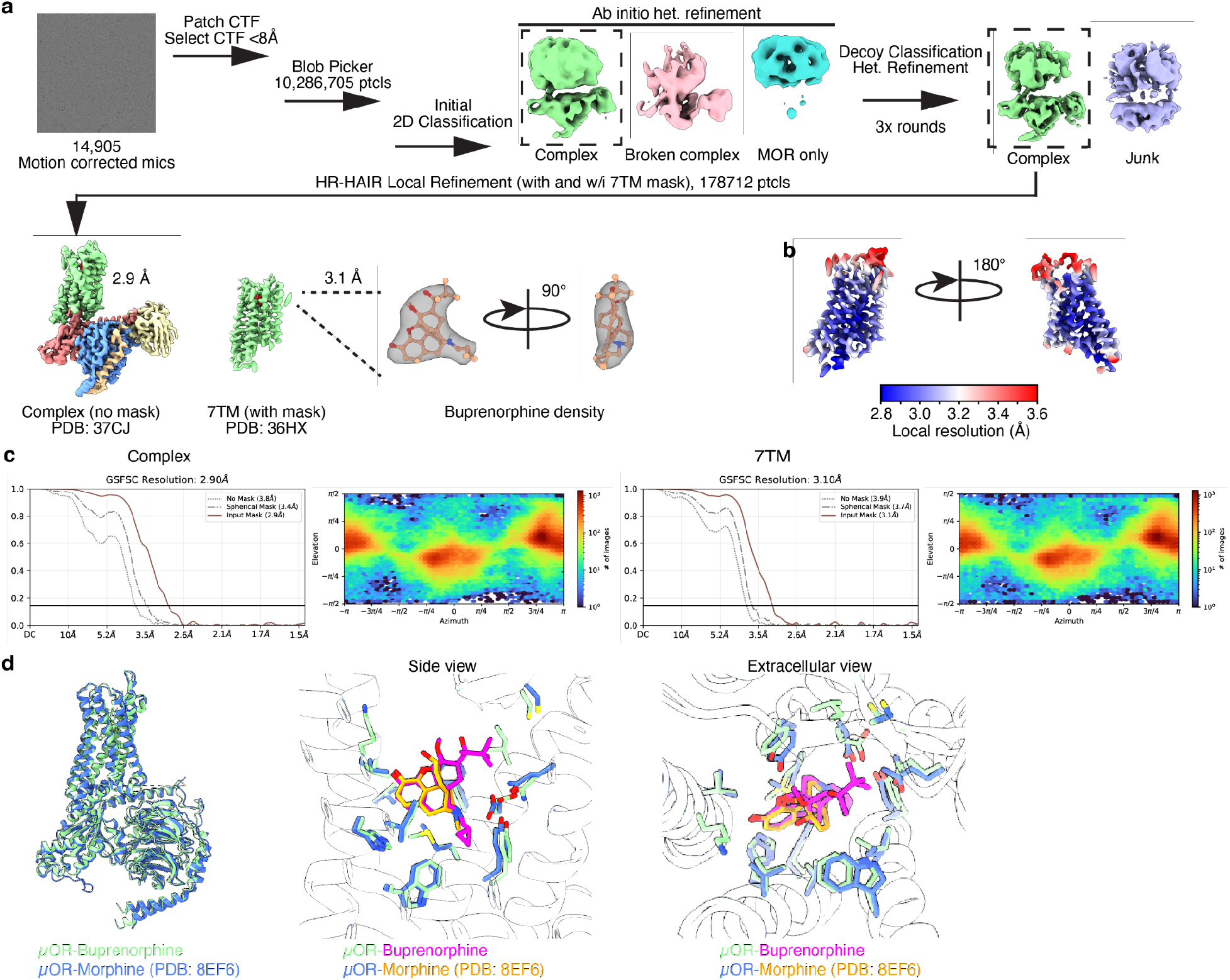
CryoEM data processing of µOR bound to Buprenorphine in complex with miniGo trimeric protein and scFv16. (a) Workflow of cryoEM data processing; motion corrected micrographs were subject to Patch CTF estimation, manual curation, and particle picking using blob picker. Particles underwent 2D classification, ab-initio reconstruction, and heterogenous refinement to obtain an initial map of the complex. This volume was used in a heterogenous refinement job using decoy volumes for junk removal. After several rounds of decoy classification, a clean particle set was subject to high-resolution ab-initio reconstruction (HR-HAIR). The resulting best class was subject to local refinement with and without a 7TM mask to obtain full complex (2.9 Å) and 7TM (3.1 Å) maps. While lower in overall resolution, 7TM map has higher map quality at the orthosteric site (b) Gold standard fourier shell correlation (GSFSC) plots (top) and viewing direction distributions (bottom) of the final reconstructions (c) Local resolution of the 7TM map.

**Supplemental Figure 9:**
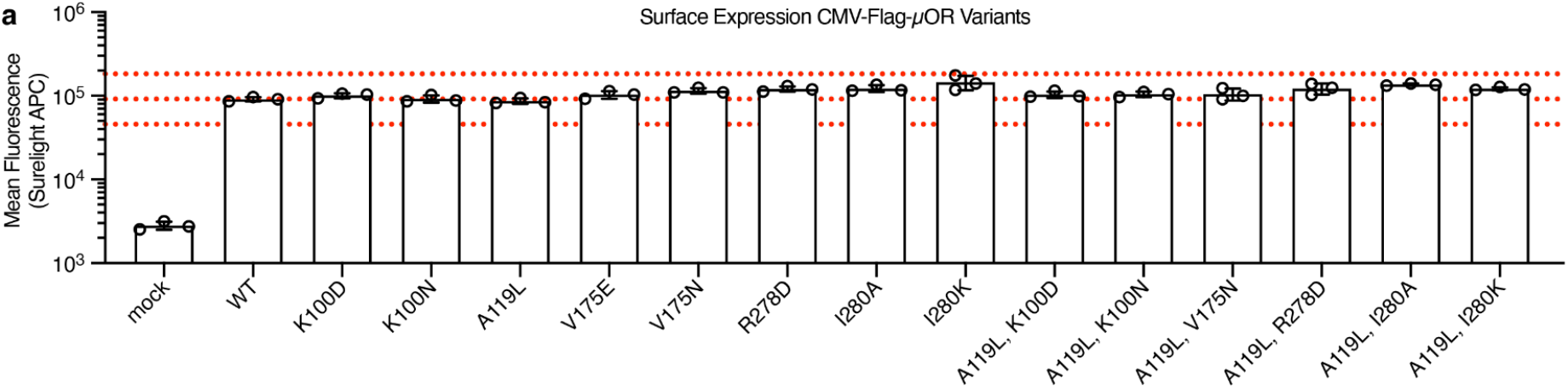
Surface Expression of Individual Mutants. (a) Relative surface expression of variants tested in this study. All are within 2 fold of WT µOR (outer red dotted lines).

**Supplemental Figure 10:**
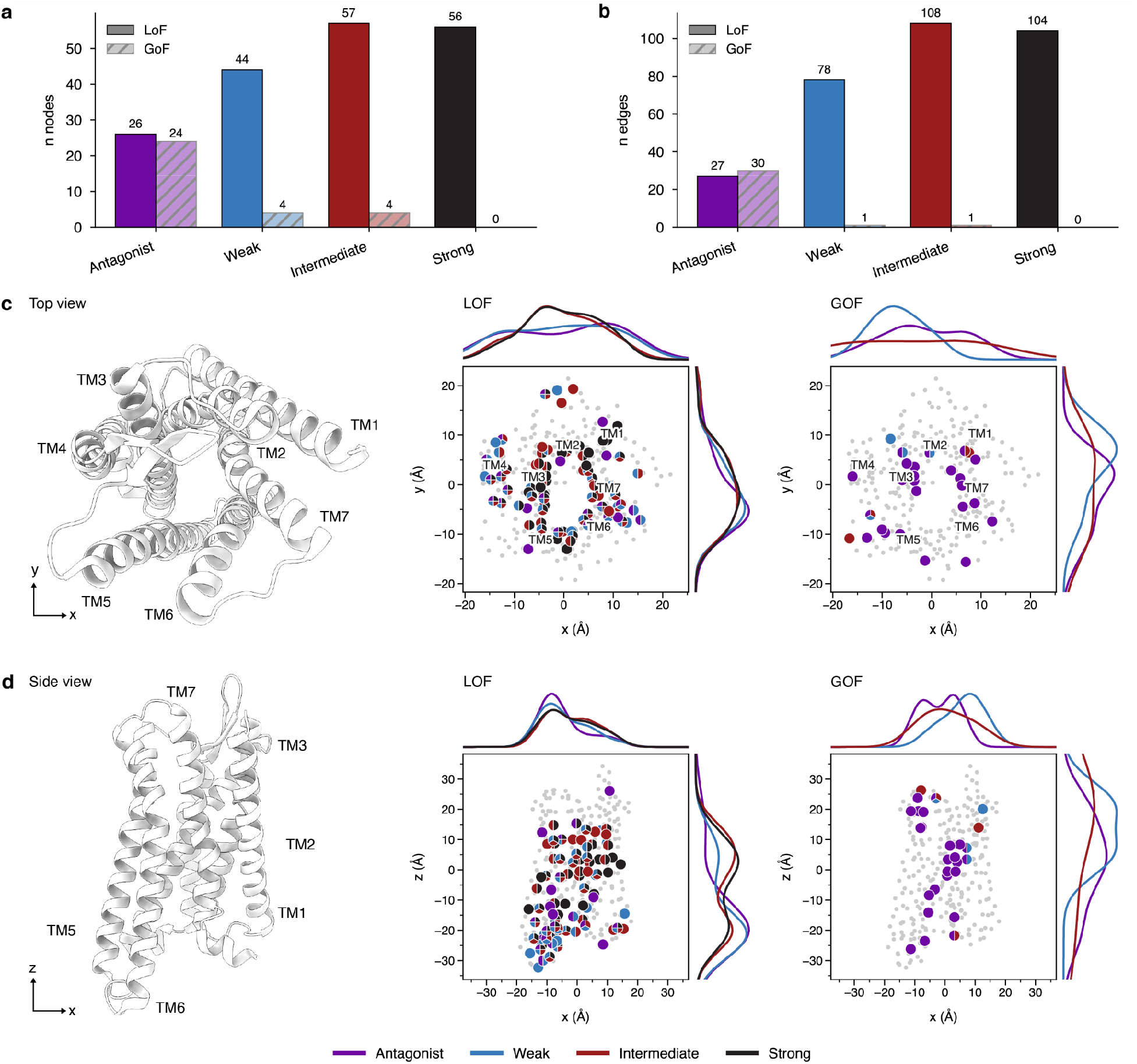
Functional residue network mapping. (a) Bar chart collating the number of nodes observed in the functional residue networks for each ligand efficacy class. (b) Bar chart collating the number of edges observed in the functional residue networks for each ligand efficacy class. (c) Top-down view of LOF and GOF nodes in each ligand efficacy class. Cartoon model of receptor shown on the left (PDB 8EFQ). Alpha carbon positions were extracted and plotted in the scatters to the right for both the loss-of-function (LOF) and gain-of-function (GOF) functional networks. Each dot represents a sigle residue and is split and colored by which ligand efficacy classes contain it as a functional node. Grey dots indicate that residue is not a functional node for any ligand efficacy class. (d) Side view of LOF and GOF nodes in each ligand efficacy class. Cartoon model of receptor shown on the left (PDB 8EFQ). Alpha carbon positions were extracted and plotted in the scatters to the right for both the loss-of-function (LOF) and gain-of-function (GOF) functional networks. Each dot represents a sigle residue and is split and colored by which ligand efficacy classes contain it as a functional node. Grey dots indicate that residue is not a functional node for any ligand efficacy class.

**Supplemental Figure 11:**
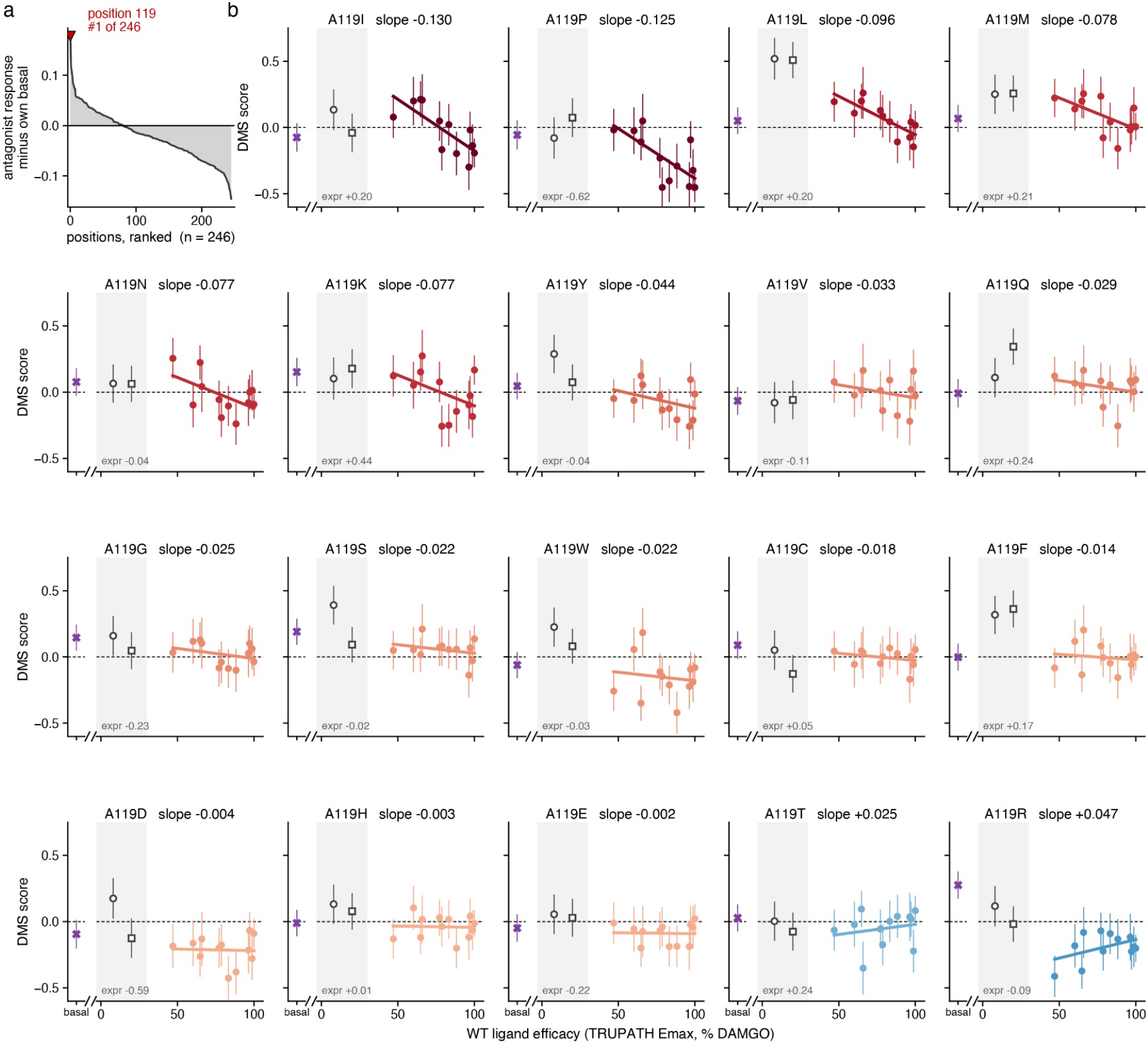
Identification of A119L change-of-efficacy variant. (a) Waterfall plot ranking receptor positions by their ability to increase signaling of antagonists when mutated. (b) Individual plots of each A119 mutation’s effect on ligand activation as measured in the mutational scanning data (y) compared to the ligands efficacy in a TRUPATH assay relative to DAMGO (x).

**Supplemental Figure 12:**
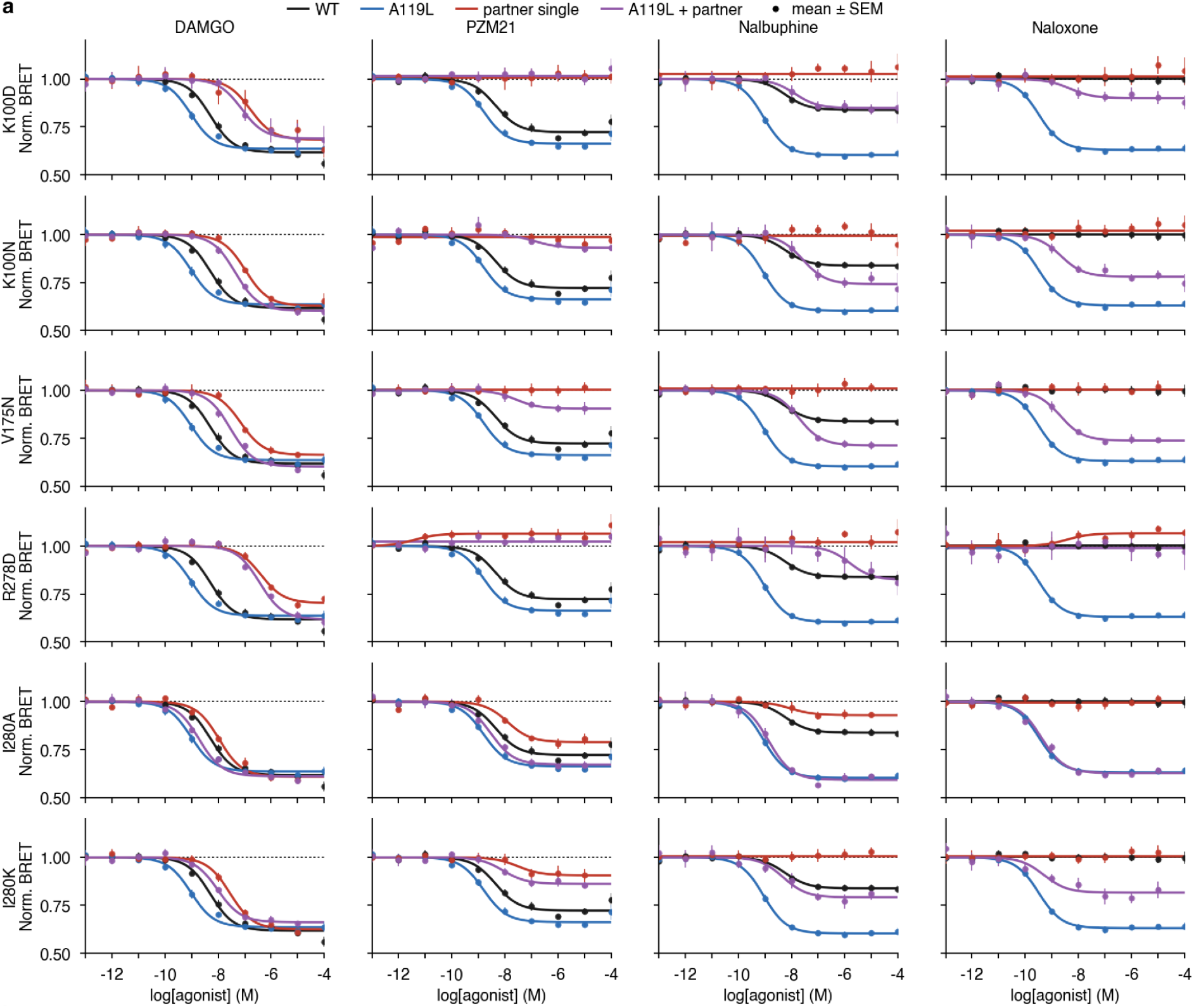
Double mutation BRET experiments. (a) Double mutant TRUPATH Gi1 BRET data testing the addition of A119L to K100D/N, V175N, R278D, and I280A/K across DAMGO, PZM21, nalbuphine, and naloxone.

## Supplemental Tables

**Supplemental Table 1:** Primer sequences for library generation Sequences of PCR primes used for generating the µOR mutational library and for amplifying the resulting library from cells after FACS screens.

**Supplemental Table 2:** Oligo sequences encoding mutations Sequences of oligonucleotides designed using the DIMPLE pipeline that encode all variants in the mutational library.

**Supplemental Table 3:** Construct sequences Sequences of plasmids used in this study. Including: HaloTag-µOR in a landing pad-compatible vector, used in mutational scanning studies, CMV-FLAG-µOR plasmid used in low-throughput signaling studies, and FLAG-MOR-miniGo112 construct for expressing µOR used in structural study.

**Supplemental Table 4: Mutational scanning scores and associated parameters.** Variant scores for each mutational scanning experiment (e.g., surface expression, concentration-response, 15-ligand screens) along with associated error metrics output from Lilace scoring pipeline. Hill and operational model parameters are included along PCA scores and external annotations (e.g., GPCRdb numbers, ligand & G protein contact distances). Position-level aggregate scores used in several plots are included. Sample data from FACS experiments is also included detailing replicates and coverage.

## Acknowledgments

We thank all members of the Coyote-Maestas and Manglik labs for their helpful feedback and discussion as we conducted this work. We acknowledge funding support from the following sources. National Institutes of Health National Institute of General Medical Sciences 5T32GM139786 (MKH), NIH National Institute on Drug Abuse DA056354 (WCM), National Institutes of Health National Institute of General Medical Sciences R35GM145238 (JSF), National Institutes of Health National Institute on Drug Abuse DA059978 (SM), National Institutes of Health National Institute of General Medical Sciences R35GM158122 (ROD), the Chan Zuckerberg Biohub (WCM and AM), Howard Hughes Medical Institute Hanna Gray Fellowship (WCM), Shurl and Kay Curci Foundation (WCM), and the Hypothesis Fund (WCM). This study was supported in part by HDFCCC Laboratory for Cell Analysis Shared Resource Facility through National Institutes of Health grant P30CA082303. We would like to thank the UCSF Cryo-EM facility staff, particularly Glenn Gilbert and David Bulkley, for training and technical assistance during data collection. Cryo-EM equipment at UCSF is partially supported by National Institutes of Health grants S10OD020054, S10OD021741 and S10OD026881 and Howard Hughes Medical Institute.

## Author contributions

MKH, WCM, and AM conceived the project. MKH, PRG, and WCM generated and cloned the µOR deep mutational library. MKH designed and performed deep mutational scan experiments with input and assistance from ZW, MVZ, AM, and WCM. JGE, BRV, and SM provided reagents. MKH, JF, JR, and CBM processed raw next-generation sequencing data with input from HP. MKH, EJF, DR, ZZ, and WCM analyzed DMS datasets with input from JF, ROD, AM. KH and KS expressed and purified the µOR construct for structural studies. MKH and KS performed cryo-EM data collection and data processing with help from cryo-EM facilities. KS and AM built and refined models of the µOR-buprenorphine structure. MKH generated receptor constructs, performed signaling studies, and analyzed the data. MKH prepared figures. MKH, WCM, and AM wrote the manuscript, with edits and approval from all authors. WCM and AM supervised the overall project.

## Data availability

Variant count data, scores, and scripts used to analyze data and prepare figures have been deposited on Github: https://github.com/matthewkarlhoward/HaloTag-MOR_DMS_analysis and Zenodo: https://doi.org/10.5281/zenodo.22051140. DMS scores and associated calculations are also included in **Supplemental Table 4**. A Shiny app has been generated for exploration and visualization of the mutational scanning datasets: https://matthewkhoward.com/mor-dms/

## Materials and methods

### MOR deep mutational library generation

The DIMPLE platform was used to generate the MOR deep mutational library ^46^. Briefly, we designed the library to contain all missense mutations at each position in MOR. We additionally included synonymous mutations and insertions and deletions of 1, 2, and 3 amino acids at each position. Insertions coded for G, GS, and GSG. These mutations were encoded in oligos with flanking BsaI sites and then ordered as a SurePrint Oligonucleotide library (Agilent Technologies) (**Supplemental Table 2**). DNA was resuspended, and the sublibrary fragments were amplified using PrimeStar GXL DNA polymerase and fragment-specific primers (**Supplemental Table 1**). These reactions were subjected to PCR cleanup using Zymo Clean and Concentrate-5 kits. The cDNA sequence of HaloTag-MOR WT was synthesized by Twist Bioscience in their High Copy Number Kanamycin backbone; BsmBI and BsaI cut sites were removed via silent mutations. For each library fragment, this plasmid was amplified to add BsaI sites, gel purified, and the corresponding oligo sublibrary was assembled using BsaI-mediated Golden Gate assembly. These reactions were cleaned using and electroporated into MegaX DH10B cells, recovered for 1 hour, and added to 30 mL LB + Kanamycin and grown while shaking at 37 C until they reached OD 0.6-0.7. DNA was isolated using a Zymo Zyppy Plasmid Miniprep kit. Each sublibrary was quantified using Invitrogen Qubit dsDNA HS assay kit and pooled in equimolar ratios. This pooled library was then assembled into a BxBI landing pad-compatible cAMP reporter vector containing a GSGSGS-P2A-PuroR cassette for positive selection and a CRE-GFP reporter ^32^.The HaloTag-MOR WT plasmid sequence is provided in **Supplemental Table 3**.

### MOR mutational library cell line generation

The HEK 293T LLP-iCasp9 cells used in this study were a gift from Doug Fowler (UW) ^47^. Cell lines for MOR WT and the MOR DMS library were generated as follows. 1ug of DNA was cotransfected with 1ug BxB1 recombinase (pCAG-NLS-BxB1, Addgene #51271) using 3.75uL lipofectamine 3000 and 5uL P3000 reagent in 6 wells of a 6 well plate. For MOR WT, 2 wells were transfected and pooled following selection. For the MOR DMS library, 18 wells were transfected in parallel. Cells were cultured in “D10” media (DMEM, 10% dialyzed FBS, 1% sodium pyruvate, and 1% penicillin/streptomycin) inside humidified incubators at 37C and 5% CO_2_. The landing pad in the cell line contains a Tet-on promoter upstream of the BxB1 recombination site and a split rapamycin analog inducible dimerizable Casp-9. Two days after transfection, we induce with doxycycline hyclate (2 µg/mL) and treat with 10 nM AP1903. Recombined cells have shifted the iCasp-9 cassette out of frame, while unrecombined cells will express the cassette and, upon treatment with AP1903, die from iCasp-9-induced apoptosis. Cells were selected for 2 days in AP1903, after which they were transitioned back to D10 supplemented with doxycycline. After two days of recovery, cells were transitioned to D10 supplemented with both doxycycline and puromycin to select for cells that have proper in-frame, full-length assemblies. Following puromycin selection for two days, cells were transitioned to D10 and expanded before freezing down or using in subsequent assays.

### Fluorescence-activated cell sorting

For flow-based assays and cell sorting, frozen stocks of cells were thawed and allowed to recover for several days in D10 media. 48 hours prior to starting the experiment, cells were split into an appropriately sized dish such that they reach ∼75% confluency by the start of the sort. 36 hours prior to starting the assay, cells were induced with doxycycline hyclate (2 µg/mL). Doxycycline was subsequently washed out after 24 hours, and cells were maintained in D10 for the remaining 12 hours prior to sorting.

### µOR surface expression deep mutational scan

The surface expression sort assays were performed as follows. Following induction of expression using doxycycline, plates were transferred to 4 °C for 10 minutes to halt receptor trafficking. Subsequently, all cells were treated with a HaloTag dye, JF635i, dissolved in D10 to a final concentration of 200 nM for 30 minutes at 4 °C. We thank Luke Lavis (HHMI Janelia) for graciously sharing this reagent. JF635i is a cell-impermeable dye that therefore conjugates only to cell surface-expressed receptors. The JF635i dye solution was aspirated. D10 (no phenol red) was then added, and cells were incubated for 30 minutes further to facilitate the washout of excess dye. Finally, cells were washed 3x with PBS-EDTA, detached using TrypLE Express for 10 minutes at room temperature, and resuspended in FACS buffer (DPBS + 3% FBS) and kept on ice prior to sorting. Cell sorting was performed using either a Beckman-Coulter Cytoflex SRT or BD FACSMelody. An example gating strategy is shown in **Supplemental Figure 2**. Briefly, cells were gated on FSC-A and SSC-A for HEK293T cells, then FSC-A and FSC-H for singlets. For the surface expression assay, we assessed expression on the APC-A channel to monitor JF635i fluorescence. For each replicate, the cell population was divided into four equal bins, each containing 25% of the population. For sorting experiments, we aimed to collect cells equal to or greater than 200x the expected number of variants in our library to ensure sufficient coverage of all variants.

### µOR Gi signaling deep mutational scans

Opioid ligands, Forskolin (Sigma), and Trimethoprim (Sigma) were diluted from 10 µM stocks (in DMSO) into D10 media containing 1 µM forskolin and the indicated ligand concentration. Media was aspirated from plates, and these ligand mixtures were applied for a total of 5h. After incubation, cells were detached using TrpyLE Express, washed, and resuspended in FACS buffer (2% FBS in TBS or DPBS). Cells were kept on ice prior to sorting. Depending on instrument availability, cell sorting was performed using a Beckman-Coulter Cytoflex SRT, BD FACSMelody, or a BD S8 cell sorter. An example gating strategy is shown in **Supplemental Figure 2**. Briefly, cells were gated on FSC-A and SSC-A for HEK293T cells, then FSC-A and FSC-H for singlets. BFP-cells were then selected for further analysis. For the Gi-cAMP assay, we assessed activity using eGFP on the FITC-A channel. For each replicate, we determined eGFP gates using the 1 µM Forskolin condition. This population was divided into four bins, each with 25% of the population. These gates were maintained for all sorts that day with DAMGO or other indicated ligands. For the concentration-response DMS experiments, all concentrations were run each day for a given ligand, e.g. DAMGO, fentanyl, or morphine. For the screens at saturation concentrations across 15 ligands, the set was split so 7 or 8 ligands were run each day due to timing constraints. Each opioid ligand was used at 10 µM. In each experiment, we aimed to collect cells equal to or greater than 200x the expected number of variants in our library.

### Genomic DNA extraction and sequencing

Following cell sorting, genomic DNA (gDNA) was extracted from cells using Quick-DNA Zymo Miniprep Plus kits. All resultant gDNA was used as a template for PCR to generate amplicons of the target gene using cell_line_for_5 and P2A_cell_line_rev primers (**Supplemental Table 1**). Some experiments used QC_for instead of cell_line_for_5 to amplify the target gene (**Supplemental Table 1**). PCR reactions were then cleaned using Zymo DNA Clean and Concentrator-25 kits, mixed with NEB Purple Loading dye (6x, no SDS), and run on a 1% agarose 1x TBE gel. Target amplicons were excised and purified using Zymo Gel DNA Recovery kits. Amplicon DNA concentrations were then quantified using an Invitrogen Qubit dsDNA HS assay kit. Libraries were prepared for deep sequencing using the Illumina Nextera XT DNA Library prep kit. Libraries were indexed using the IDT for Nextera Unique Dual Indexes or Illumina Unique Dual indexes. Lengths of indexed libraries were quantified using the Agilent TapeStation HS D5000 assay, and concentrations were determined using the Invitrogen Qubit dsDNA HS assay kit. Samples were pooled and then paired-end sequenced (PE150) using either an Illumina NovoSeq 6000 or Novoseq X.

### Next-generation sequencing data processing

Sequencing files were obtained from the sequencing core as fastq.gz after demultiplexing. The experiment was processed using a DMS-specific pipeline we developed, “Dumpling”, released as part of Rosace ^75^. Dumpling implements the following steps: first, adapter sequences and contaminants were removed using BBDuk, then paired reads were error corrected with BBMerge and mapped to the reference sequence using BBMap with 15-mers (all from BBTools ^76^). Variants in the mapped SAM file were called using the AnalyzeSaturationMutagenesis tool in GATK v4 ^77^. The output of this tool is a CSV containing the genotype of each distinct variant and the total number of reads for each sample. This was then further processed using a Python script which filtered out sequences that were not part of the designed variants.

### Variant scoring and concentration–response curve fitting

Variant scores were calculated with Lilace, a statistical model for scoring variants from FACS-based deep mutational scanning data ^78^. For the surface-expression and 15-ligand saturating-concentration screens, Lilace was run with default settings where the within-screen synonymous variants are used as a baseline. Scores were centered at zero for synonymous variants; gain-of-function variants (increased fluorescent signal relative to synonymous) were positive and loss-of-function variants (decreased signal) were negative. For the concentration-response experiments the synonymous variants in the no ligand condition were used as a baseline.

Per-variant concentration–response data were fit with a three-parameter Hill equation E(x) = E_min_ + (E_max_ − E_min_) / (1 + 10^(logEC_50_ − x) with x = log₁₀[ligand] and the Hill coefficient fixed at 1, by bounded nonlinear least squares (SciPy curve_fit). Morphine and fentanyl were titrated over eight concentrations (−11.5 to −4.5 and −12.0 to −5.0 log₁₀ M) and DAMGO over six (−5 to −9 log₁₀ M, with the ligand-free condition included as a baseline anchor at −12). Fits required at least four measured concentrations, with E_min_ and E_max_ bounded to [−2, 2] and logEC_50_ to the assayed range. To reduce the influence of aberrant points, each curve was fit once, points deviating by more than twice the root-mean-square residual were removed (at most two per variant), and the curve was refit. E_min_, E_max_, and EC_50_ were taken from the refit. Only well-constrained fits were retained for analysis — those reaching a lower and an upper asymptote within the tested range, so that E_max_ and EC_50_ are determined by the data rather than extrapolated; fits that were flat or did not define both asymptotes were excluded. E_max_ was placed on an activity scale normalizing each ligand’s synonymous variants (mean synonymous E_min_ = 0, E_max_ = 1), and EC_50_ is reported as log₁₀ M.

### Operational model fitting

Each variant was fitted with a dose-response curve with the operational model of agonism ^4^, a pharmacological equation whose τ parameter is an efficacy measure that accounts for receptor expression. We combined τ with measurements of cell surface expression for each variant to split the efficacy change into an expression and coupling component. For each variant we used its response measured in the dose-response deep-mutational scan. Each concentration also carried a measurement uncertainty (a standard deviation), which we used to weight the fit so that noisier points count less. How much receptor reaches the cell surface was measured separately. For one agonist, the operational model predicts the response as a function of concentration.

Using [A] as the concentration of agonist, the response([A]) = E_m_ · τ·[A] / (K_A_ + (1 + τ)·[A]), and the observed_score_([A]) = B − response([A]). The fitted quantities are: E_m_, the largest response the cellular system can physically produce (defined here as the 99.5th percentile of the strongest response across all variants); τ, the efficacy/reserve parameter; K_A_, the concentration at which the ligand half-occupies the receptor; and B, the flat baseline of the curve. We fixed the model’s “steepness” exponent at 1. The following relationships arise: E_max_ = E_m_ · τ / (1 + τ), which rearranges to τ = E_max_ / (E_m_ − E_max_), and EC₅₀ = K_A_ / (1 + τ), where EC₅₀ is the concentration giving a half-maximal response (a potency measure)

Each variant curve was using robust nonlinear regression. Each data point was weighted by its inverse measurement uncertainty. Starting values for the fit were derived from simple features of the raw curve (its plateau, its drop, and its midpoint) so the optimizer began near a sensible solution. We estimated the uncertainty of each fitted parameter from the standard statistical approximation based on the fit’s local curvature. A curve was flagged “saturating” when its response reached ≥ 97% of the ceiling E_m_, meaning efficacy could no longer be pinned down precisely. Fits were run for all variants (10,106 morphine variants converged).

The number of working surface receptors scales in proportion to how much protein reaches the surface, a variant’s efficacy thus separates into two factors: τ*_variant_ = τ*_syn_ × S × ρ where τ_syn_ is the wild-type receptor’s efficacy, S is the variant’s surface expression relative to normal, and ρ is the per-receptor coupling efficiency (ρ = 1 for wild type). Taking logarithms turns the product into a subtraction: log₁₀ ρ = (log₁₀ τ*_variant_ − log₁₀ τ*_WT_) − log₁₀ S. We set the wild-type reference τ*_syn_* to the median efficacy of well-fit synonymous variants. ρ is the quantity of interest because it distinguishes two situations that look identical in a raw readout: a variant that signals weakly because too little of it reaches the surface (low S, normal ρ) versus one that reaches the surface normally but each receptor couples worse (normal S, low ρ). We estimated surface expression S using the fluorescence fold-change from the JF635i labeling and cell sorting experiment. Cells were stained for surface receptor and sorted by a FACS into 4 equal-sized bins ordered from lowest to highest surface fluorescence. The cells in each bin were then DNA-sequenced to count every variant. For each variant we converted its counts into the fraction of its cells falling in each bin, and computed a mean surface brightness by weighting each bin by a representative fluorescence value read from the sort histograms. Expression relative to wild type was then the difference in log-brightness. Following this, the decomposed values for each parameter were extracted and plotted.

### Ligand reweighting analysis

Morphine and fentanyl mutational effects were compared using the parameters from the three-parameter fits. Per-position effects were the mean effect across missense variants at each position (sigmoid fits only, ≥ 5 missense variants per position), expressed relative to that ligand’s synonymous mean: right-shifted log₁₀ EC_50_ for potency and reduced activity-scaled E_max_ for efficacy. A position was defined as loss-of-function for a ligand if its effect exceeded one standard deviation of that ligand’s synonymous distribution.

The overall similarity of the two ligands’ landscapes was quantified by Spearman correlation of per-position effects (E_max_ ρ = 0.91; EC_50_ ρ = 0.66; corresponding variant-level correlations ρ = 0.76 and 0.34). To determine whether the two ligands diverge by more than experimental noise, we established a within-ligand reproducibility ceiling by bootstrap resampling missense variants within each position (10,000 iterations, fixed seed): for each iteration, the Spearman correlation between two independent resamples of the same ligand gave the highest correlation attainable if the two ligands were identically weighted. For both properties the cross-ligand correlation fell below this ceiling (E_max_, cross 0.86 vs ceiling 0.93– 0.96; EC_50_, cross 0.57 vs ceiling 0.82–0.86; 100% of iterations below ceiling). Individual reweighted positions were identified from the bootstrap distribution of the per-position morphine − fentanyl difference, tested for departure from zero and corrected across positions that were loss-of-function for at least one ligand by the Benjamini–Hochberg procedure (q < 0.05 significant). To establish that reweighting was distributed rather than driven by these positions, the reproducibility-ceiling comparison was repeated after excluding them; for Emax the cross-ligand correlation remained below the ceiling (gap 0.097 → 0.087, 100% of iterations), indicating divergence spread across many individually sub-threshold positions.

Mutations shifted morphine EC_50_ more than fentanyl across the whole receptor, so the per-position EC_50_ difference was centered on its across-position median before significance testing and structural coloring (**Figure 4C**), so that it reflects position-specific reweighting rather than this receptor-wide difference. The E_max_ difference showed no comparable offset and was analyzed and displayed without centering (**Figure 4F**). Structures were colored on a diverging scale by the per-position morphine − fentanyl difference (morphine, blue; fentanyl, red).

### Determination of primary and second shell ligand contacts

Structures for fentanyl (8EF5), morphine (8EF6), mitragynine pseudoindoxyl (MP, 7T2G), naloxone (9PXY), TRV130 (oliceridine, 8EFB), PZM21 (8EFO), C6-Guano (7U2K), SR17018 (8EFL), DAMGO (8EFQ), lofentanil (7T2H) were obtained from the PDB. Ligands and receptor chains were identified. A structure of buprenorphine bound to µOR was determined as described herein. Methadone, naltrexone, nalbuphine, and butorphanol poses were determined using Chai-1. Specifically, the human µOR sequence was input along with the SMILES string for each ligand. For methadone, nalbuphine, and butorphanol, the sequence of GNAI was included to induce an active state of the receptor. For naltrexone, no G protein was included. All predictions were setup using MSA. Ligand poses were nearly identical among the five predictions generated for each ligand. For experimental and predicted poses, first shell residues were defined as those with a heavy atom within 4.5 Å of any ligand heavy atom. Second shell residues were defined as those within 5 Å of any first shell residue. All other receptor residues were not categorized.

### Hierarchical clustering and principal component analysis of multi-ligand dataset

All analyses were restricted to missense variants at positions 65–355 (the transmembrane region) to filter out the low-signal N- and C-terminal tails. This is the same position window used for the stacked heatmap in **Figure 6B**. Variants were required to have complete effect measurements across all 15 ligands (no missing values), yielding 5,519 missense variants. For position-level analyses these were collapsed to a per-position mean missense effect for each ligand, giving 291 positions with complete data across all 15 ligands.

For the clustering analysis, each ligand was represented by its profile of per-position mean missense effects over positions 65–355 (positions present for all ligands). Pairwise distances between ligand profiles were computed as correlation distance, 1 − Pearson r, so that ligands are grouped by profile *shape* — which positions are mutation-sensitive — independently of overall effect magnitude. Ligands were then clustered by Ward’s minimum-variance linkage (ward.D2). The resulting dendrogram was rotated toward the measured Emax efficacy ordering — a topology-preserving rotation with no branch crossings, so the displayed leaf order reflects genuine clustering structure rather than an imposed order. DAMGO was pinned to the bottom row. FSK (No Ligand) was not clustered and is shown pinned to the top of the heatmap.

PCA was performed on the same missense-effect matrix. Each column is one ligand; each row is either a single variant (variant-level PCA; 5,519 × 15) or a single position (position-level PCA on per-position mean effects; 291 × 15). Each ligand (column) was standardized to zero mean and unit variance (z-scored), the matrix was mean-centered, and principal components were obtained by singular value decomposition. Per-row PC coordinates (positions/variants) are referred to as scores and per-ligand weights (V·S) as loadings. The variant-level PCA was sign-aligned to the position-level PCA (matching the per-position mean of each PC) so the two share a canonical orientation; downstream sign is therefore consistent across panels.

### Functional residue network generation

Networks were built from the per-variant DMS effect scores, restricted to the transmembrane region. For each of the 15 ligands, the distribution of synonymous variants defined a per-ligand null: mean and standard deviation of the synonymous effect. A missense variant was scored as loss-of-function (LOF) or gain-of-function (GOF) if it exceeded the synonymous mean by > 2 SD in the positive or negative direction. To remove variants whose apparent signaling change is confounded by folding/trafficking loss, only variants with near-wild-type surface expression were counted (surface expression score ≥ (synonymous surface mean − 2 SD). Variants below this cutoff were excluded before any counting. Per-position disruption count (“severity”) were computed as the number of distinct amino-acid substitutions at that position that crossed the 2-SD LOF (or GOF) threshold, among the surface-passing variants. Ligands were grouped into efficacy classes determined by their clustering and efficacy as in **Figure 5-6** and a class-level per-position count was taken as the mean disruption count across the class’s ligands. This class disruption count is the value driving node severity/size displayed in the figure. Within each class and direction (LOF for panel A, GOF for panel B), a position was promoted to a network node if it satisfied both criteria: class disruption count ≥ max(2, 75th percentile of that class’s non-zero counts). The Q75-of-nonzero cut adapts the stringency to each class, with a hard floor of 2 substitutions. Nodes were placed at their Cα positions in the active-state µ-opioid receptor structure (PDB 8EFQ). An edge was drawn between any two nodes whose Cα–Cα distance was < 8.0 Å. Edges therefore mark spatial proximity among disruption-sensitive residues, revealing physically contiguous functional clusters. Each panel was rendered in ChimeraX as a translucent receptor cartoon with opaque node spheres and edge cylinders. Node sphere radius encodes severity: it is area-proportional to the class disruption count (radius ∝ √count), range-normalized within each direction onto 0.55–2.40 Å — so LOF and GOF panels each span the full size range but a LOF and a GOF sphere of equal radius do not necessarily represent the same absolute count.

### cAMP inhibition flow cytometry assays

Cells expressing HaloTag-MOR WT were cultured and induced with doxycycline as described above for the mutational scanning experiments. Here, ligands were tested in concentration-response on the WT receptor to determine EC_50_ and E_max_ values for each ligand in the assay. After ligand incubation time of 5 hours, cells were detached, washed in DPBS with 2% FBS, then assayed using an Agilent Quanteon flow cytometer. Gating was similar to for the mutational scanning experiments: gate for HEK293T cells using FSC and SSC, then signets using FSC-A and FSC-H, BFP-, then eGPF intensity using the FITC channel. Here, each ligand x concentration eGFP signal was calculated as a ratio relative to forskolin only. Curves were plotted in Prism and parameters derived from fitting a 3-paramater Hill equation to the datasets.

### TRUPATH BRET Assays

TRUPATH BRET assays to monitor G protein dissociation were performed using established protocols ^79^. The sequence of wildtype human µOR with an HA signal peptide and FLAG tag appended on the N-terminus was synthesized in a pTwist-CMV vector (Twist Biosciences). Mutational variants of the µOR were generated using a Q5 Site-Directed Mutagenesis Kit (NEB) following manufacturers’ protocol. HEK293T cell lines (ATCC) were cultures in DMEM (Gibco) supplemented with 10% dialyzed FBS (Gibco). Cells were plated in a 6-well dish at appropriate density to reach ∼80% confluent at the time of transfection. Cells were transfected using Lipofectamine 3000 (Invitrogen) following manufacturers’ protocol using a 1:1:1:1 ratio of µOR:Gαi1:Gβ3:Gγ9 constructs (total of 2 µg DNA) per well. After 12 hours, cells were lifted using TrypLE Express (Gibco), counted, and plated in pre-coated poly-lysine white 384 well plates at a density of ∼20,000 cells in DMEM supplemented with 1% dialyzed FBS. Cells were allowed to attach overnight (12 hours). Assay buffer was prepared by mixing Prolume Purple 1 mM (Nanolight Technology) with HBSS (Gibco) to a final concentration of 7.5 µM. The growth medium was then removed by gently inverting and shaking the plate and blotting dry. Assay buffer was then aliquoted, 20 µL into each well. 10 µL prepared ligand dilution solutions (prepared in HBSS) were allotted into each well. SR-17018 precipitated in HBSS, PBS, and each buffer supplemented with 1% tween 80, so the data reported in **Figure 5C** is a single concentration (100 µM) as a reliable curve was not obtained. The plate was then loaded into a Clariostar (BMG Labtech) platereader and read for four cycles and the fourth cycle was used for all analysis. BRET ratios were computed as the ratio of GFP2 emission to RLuc8 emission (515/410). Data were plotted in Prism.

### Expression testing of µOR variants used in TRUPATH assays

Cell surface expression of pTwist-CMV-µOR variants was measured via flow cytometry. Plasmids were transfected into HEK293T using Lipofectamine 3000 (Invitrogen) as described above. After 24 hours, cells were lifted using TrypLE Express (GIBCO), washed twice with DPBS + 3% dialyzed FBS. Cells were then incubated with SureLight APC Anti-DDDDK tag antibody [M2] (Abcam) diluted 1:200 in DPBS + 3% dialyzed FBS for 30 minutes in the dark. Following this, cells were washed twice with DPBS + 3% dialyzed FBS and analyzed using flow cytometry, monitoring cell surface expression using the APC channel. All variants were found to express within 2-fold of WT µOR.

### Ligands

**Ligands:**
| Compound | Vendor / cat# |
| --- | --- |
| Carfentanil | Cayman Chemical Cat # 19884 |
| Butorphanol Tartrate | Cayman Chemical Cat # 28625 |
| Buprenorphine Hydrochloride | Sigma Aldrich Cat # PHR1729 |
| SR17018 | Cayman Chemical Cat # 24480 |
| DAMGO | Tocris Cat # 1171 |
| Fentanyl Citrate | Spectrum Cat # F1147 |
| Morphine Sulfate | Spectrum Cat # M1167 |
| Naloxone Hydrochloride | Tocris Cat # 0599 |
| Naltrexone Hydrochloride | Sigma-Aldrich Cat # N3136 |
| Mitragynine Pseudoindoxyl | Cayman Chemical Cat # 20033 |
| Nalbuphine Hydrochloride | Sigma-Aldrich Cat # N-051 |
| PZM21 | Cayman Chemical Cat # 20576 |
| TRV130 (Oliceridine) | Cayman Chemical Cat # 30624 |
| C6-Guano | Majumdar lab, Washington University in St. Louis |
| Methadone Hydrochloride | Spectrum Cat # M1426 |
| Trimethoprim | Thermo Scientific Chemicals Cat # AC455120050 |

### Expression and Purification of MOR-miniG_o1_12

The human wildtype MOR (Uniprot: P35372) was cloned into pCDNA-Zeo-TetO, a custom pcDNA3.1 vector containing a tetracycline-inducible gene expression cassette ^80^. The resulting MOR construct comprised an N-terminal influenza haemagglutinin signal sequence followed by an M1-FLAG (DYKDDDD) epitope tag. The C-terminus of MOR was fused to the miniG_o1_12 protein via a linker sequence containing a human rhinovirus 3C (HRV 3C) protease cleavage site and a Go N-terminal fragment to enable scFv16 engagement. The resulting MOR-miniG_o1_12 was transiently transfected into inducible Expi293F TetR cells (ThermoFisher Scientific) using FectoPRO transfection kit (Polyplus-Sartorius) following manufacturer instructions. The cells were untested for mycoplasma contamination and were cultured in a shaking incubator maintained at 37°C and 8% CO_2_. After 16 hrs, protein expression was induced with 1 μg/mL doxycycline hyclate (Sigma-Aldrich). Cells were harvested 48 hrs post induction by centrifugation at 10,000 x g for 15 mins. Cell pellets were flash frozen in liquid N_2_ and stored at −80°C.

All protein purification steps were done at 4°C. Thawed cell pellets from a 250 mL Expi293 expression harvest were resuspended in 50 ml hypotonic lysis buffer comprising 20 mM HEPES (pH 7.5), 1 mM EDTA, a Pierce^TM^ Protease Inhibitor Tablet, and 50 nM Buprenorphine (Bup; Spectrum). Cells were lysed by stirring for 15 min at 4°C and harvested by centrifugation at 38,000 x g for 20 mins. Cell pellet was dounce-homogenized in solubilization buffer comprising 20 mM HEPES, pH 7.5, 150 mM NaCl, 1% (w/v) n-Dodecyl-β-D-Maltopyranoside (DDM; Anatrace), 0.1% (w/v) cholesteryl hemisuccinate (CHS, Sigma-Aldrich), 0.3% CHAPS, 2 mM MgCl_2_, 2 mM CaCl_2_, 5 mM adenosine 5′-triphosphate (ATP; Fisher Scientific), 100 μM TCEP, 50 nM Bup, 3 ul benzonase (MilliporeSigma), and a Pierce^TM^ protease inhibitor tablet. The sample was stirred for 1 h, and the detergent-solubilized fraction was clarified by centrifugation at 50,000 x g for 30 min. The detergent-solubilized sample was supplemented with 2 mM CaCl_2_ and incubated in batch with 1 mL homemade M1-FLAG-antibody conjugated CNBr-Sepharose under slow rotation for 1 hr. The resin was transferred to a glass column and washed with 50 mL of ATP wash buffer comprising 20 mM HEPES, pH 7.5, 150 mM NaCl, 0.1% (w/v) DDM, 0.01% (w/v) CHS, 0.03% CHAPS, 5 mM ATP, 2 mM CaCl_2_, 2 mM MgCl_2_, 100 μM TCEP, 50 nM Bup. This was followed by 20 mL of DDM Wash buffer comprising 20 mM HEPES, pH 7.5, 150 mM NaCl, 0.1% (w/v) DDM, 0.01% (w/v) CHS, 0.03% CHAPS, 2 mM CaCl_2_, 100 μM TCEP, 50 nM Bup. Receptor was slow exchanged into GDN buffer by 10 mL washes containing the following ratios of DDM wash buffer to GDN wash buffer: 1:1, 1:3. 1:7. GDN Wash buffer was comprised of 20 mM HEPES, pH 7.5, 150 mM NaCl, 0.2% (w/v) glyco-diosgenin (GDN; Anatrace), 0.02% (w/v) CHS, 0.03% CHAPS, 2 mM CaCl_2_, 100 μM TCEP, 50 nM Bup. This was followed by a 20 mL wash with GDN wash buffer. Receptor-containing fractions were eluted with 20 mM HEPES, pH 7.5, 150 mM NaCl, 0.0075% (w/v) GDN, 0.00075% (w/v) CHS, 5 mM EDTA, 0.2 mg/mL FLAG peptide, 100 μM TCEP, 50 nM Bup. Fractions containing receptor were concentrated in a 50-kDa MWCO spin filter (Amicon) and subjected to size-exclusion chromatography (SEC) on a Superdex 200 Increase 10/300 GL column (Cytiva), which was equilibrated with 20 mM HEPES, pH 7.5, 150 mM NaCl, 0.0075% (w/v) GDN, 0.00075% (w/v) CHS, 100 μM TCEP, 50 nM Bup. Fractions containing monodisperse MOR-miniG_o1_12 were combined and concentrated in a 50-kDa MWCO spin filter before complexing with Gβ_1_γ_2_ and scFv16.

### Expression and Purification of G_β1γ2_ heterodimer

The human G_β1_ subunit with an N-terminal 6x His-tag, and untagged G_γ2_ were cloned into a bicistronic pVLDual vector. A baculovirus was generated in Spodoptera frugiperda Sf9 insect cells (unauthenticated and untested for mycoplasma contamination, Expression Systems). The G_β1γ2_ was produced in HighFive insect cells grown in ESF921 medium (Expression systems). Cells at a density ∼3 × 10^6^ cells/ml were infected with 10 ml/lt Gβ_1_γ_2_ baculovirus. After 48 hours of incubation, cells were harvested by centrifugation at 4000 x g for 20 mins and stored in −80°C. Cell pellets were thawed and resuspended in lysis buffer (200 ml buffer per liter of cell culture pellet) comprising of 10 mM HEPES pH 7.5, 0.1 mM MgCl2, 5 mM β-mercaptoethanol (β-ME), 10 μM guanosine diphosphate (GDP), and a Pierce^TM^ Protease Inhibitor Tablet. Cell membranes were centrifuged at 38,500 x g for 15 mins. The pellet was resuspended in solubilization buffer (20 mM HEPES pH 7.5, 100 mM NaCl, 1% sodium cholate, 0.05% DDM, 5 mM MgCl2, 2 µL CIP (New England Biolabs), 5 mM β-ME, 20 mM imidazole, 10 μM GDP), and a Pierce^TM^ Protease Inhibitor Tablet using a Dounce homogenizer (200 ml buffer per liter of cell culture pellet). Resuspended membranes were then stirred for 1 hour at 4°C. To pellet insoluble debris, the solubilized membranes were centrifuged at 38,500 x g for 20 mins. The supernatant was mixed with 8 mL Ni-NTA resin prewashed with solubilization buffer. To enable batch binding, solution was incubated for 2 hours at 4°C with gentle rotation. Ni-NTA resin was collected by centrifugation at 500 x g for 10 mins. Ni-NTA resin was washed twice with 40 ml solubilization buffer per wash. For slow exchange of protein into buffer without sodium cholate, Ni-NTA resin was transferred to a gravity-flow column and washed 5 times with 20 ml buffer each time. These washes contain the following ratios of E1 and E2 buffers: 50:50, 25:75, 10:90, 5:95, 0:100 where E1 was solubilization buffer supplemented with 5 mM Imidazole and E2 was 20mM HEPES pH 7.50, 100 mM NaCl, 0.05% DDM, 1 mM MgCl2, 5 mM β-ME, Pierce^TM^ Protease Inhibitor Tablet, and 20 mM Imidazole. A final wash in 25 ml of E2 buffer supplemented with 30 mM Imidazole was followed by elution using 40 ml of E2 buffer supplemented with 250 mM imidazole. Elution was incubated with 3 μl lambda phosphatase (supplemented with 1 mM MnCl2 for activity), 1 µL CIP (NEB), 1 uL Antarctic phosphatase, and incubated at 4°C for 30 min with gentle rocking. The protein was then dialyzed using 10 kDa cutoff snakeskine membrane against in 1 L of dialysis buffer comprising 20 mM HEPES 7.5, 100 mM NaCl, 0.05% DDM, 1% Glycerol, 100µM TCEP for 16 hours. Protein was then centrifuged at 38,500 x g for 20 mins to pellet precipitated protein. The supernatant was decanted into an equal volume of Anion Exchange buffer A comprising 20mM HEPES pH 7.50, 0.05% DDM, 100µM TCEP. Protein sample was loaded onto a 5 mL HiTrap Q FF antion exchange chromatography column (Cytiva) and eluted with a gradient of 5-50% Anion exchange buffer B comprising of 20mM HEPES pH 7.50, 1M NaCl, 0.05% DDM, 100µM TCEP over 20 column volumes. Largest peak fractions corresponding to G_β1γ2_ were pooled and concentrated using a 30 kDa Amicon spin filter. Concentrated protein was supplemented with 20% Glycerol, aliquoted, flash frozen in liquid nitrogen, and stored at −80°C.

Expression and Purification of antibody fragment scFv16. Protein scFv16 was expressed and purified as previously reported ^81^. In brief, scFv16 was expressed in Hi5 insect cells infected with baculovirus in a secreted format. Cell medium was pH balanced using Tris pH 7.5 and chelating agents were quenched by supplementing 1mM nickel chloride and 5mM calcium chloride and incubation with stirring for 1h at 25°C. Protein was purified using Ni-affinity chromatography and the His-tag was cleaved using HRV-3C protease during dialysis. Cleaved protein was further purified by reloading over Ni-NTA resin. The flow through was collected and purified over a size-exclusion chromatography using a Superdex 200 16/60 column (GE healthcare). Monomeric fractions were pooled, concentrated and flash frozen in liquid nitrogen until use.

### Complex formation of MOR-miniG_o1_12 with G_β1γ2_ heterodimer and scFv16

Purified Gβ1γ2 heterodimer and scFv16 were mixed with MOR-miniG_o1_12 in 2 molar excess. The complex was incubated at 4°C for 16 hours and then injected onto a Superdex200 Increase 10/300GL (Cytiva) gel filtration column equilibrated in 20 mM HEPES, pH 7.5, 150 mM NaCl, 0.0075% (w/v) GDN, 0.00075% (w/v) CHS, 100 μM TCEP, 50 nM Bup.) to isolate monodisperse material for cryoEM grid preparation.

### CryoEM sample preparation and data collection

UltrAuFoil (R1.2/1.3) 300 mesh grids (Quantifoil) were glow discharged and 2.5 ul of the MOR-miniG_o1_12-Gβ1γ2-scFv16 complex with Buprenorphine was applied to the grids before plunge freezing in liquid ethane using a Vitrobot Mark IV (ThermoFisher) using 2-8 s blot time, 10 s wait time, 0 blot force while maintaining 4°C temperature and 100% humidity. Movies were recorded using EPU (ThermoFisher) on a Titan Krios G4 (ThermoFisher) equipped with a TFS SelectrisX energy filter and a TFS Falcon 4i direct electron detector. A 165,000 x nominal magnification and a pixel size of 0.7262 Å/pixel, was used with a dose of 45.51 e-Å^-2^.

### CryoEM data processing, model building, and refinement

Raw movies were imported into cryoSPARC v4.5.3 (Structura Biotechnology Inc) and motion correction using Patch Motion Correction. Contrast transfer functions were calculated using Patch CTF estimation. A threshold of CTF fit resolution higher than 8 Å was used to exclude low-quality micrographs. Particles were picked using blob picker with a particle diameter range from 70 Å to 150 Å, and picked particles (n= 7,193,094) were extracted with a box size of 288 pixels binned to 72 pixels. An ab-initio model was generated and used as input volume in a heterogenous refinement job with 3 additional junk collection volumes produced by early termination of an ab-initio job. This strategy, called decoy classification^82^, has worked well for GPCR datasets. Particles were cleaned up with multiple rounds of 3D classification by heterogenous refinement. Particles were subsequently unbinned and sorted by multiple rounds of ab-initio reconstructions and heterogenous refinements.

A clean particle stack (n = 381,782) was subjected to high resolution heterogeneous ab initio reconstruction (HR-HAIR ^83^) with the following parameters: no. of classes 3, initial resolution 5Å, final resolution 2.3 Å, fourier radius step 0.005, initial minibatch size 300, final minibatch size 1000. A single class of particles (n = 296,187) emerged as the desired complex and was subject to another round of HR-HAIR yielding two classes of the desired complex. Resulting volumes were subjected to homogenous reconstruction and local refinement before being manually inspected to choose the best class. This class (n= 178,712) had an overall resolution of 2.9 Å and was used to model the full complex structure. To better resolve the Buprenorphine density in the orthosteric site, local refinement using a focus mask that covers the seven transmembrane domain of MOR yielded a 3.1 Å reconstruction. Model building and refinement were carried out using PDB 8EF6 as a starting model, which was fit into the cryoEM map using ChimeraX ^84^. Subsequent rounds of refinement were carried out using real space refinement in Phenix ^85^ with manual adjustments in Coot ^86^. The Buprenorphine model and restraints were generated with eLBOW and docked using Coot. The final structure for the full complex and the locally refined 7TM domain have been deposited in the PDB with accession codes 37CJ and 36HX respectively.

## References

1. Ahlquist, R. P. A study of the adrenotropic receptors. Am. J. Physiol. 153, 586–600 (1948).

2. Kenakin, T. Efficacy at G-protein-coupled receptors. Nat. Rev. Drug Discov. 1, 103–110 (2002).

3. Stephenson, R. P. A modification of receptor theory. 1956. Br. J. Pharmacol. 120, 106–20; discussion 103-5 (1997).

4. Black, J. W. & Leff, P. Operational models of pharmacological agonism. Proc. R. Soc. Lond. B Biol. Sci. 220, 141–162 (1983).

5. Rosenbaum, D. M. et al. GPCR engineering yields high-resolution structural insights into beta2-adrenergic receptor function. Science 318, 1266–1273 (2007).

6. Weis, W. I. & Kobilka, B. K. The molecular basis of G protein-coupled receptor activation. Annu. Rev. Biochem. 87, 897–919 (2018).

7. Cherezov, V. et al. High-resolution crystal structure of an engineered human beta2-adrenergic G protein-coupled receptor. Science 318, 1258–1265 (2007).

8. Warne, T. et al. The structural basis for agonist and partial agonist action on a β(1)-adrenergic receptor. Nature 469, 241–244 (2011).

9. Zhao, J. et al. Ligand efficacy modulates conformational dynamics of the µ-opioid receptor. Nature 629, 474–480 (2024).

10. Manglik, A. et al. Structural insights into the dynamic process of β2-adrenergic receptor signaling. Cell 161, 1101–1111 (2015).

11. Nygaard, R. et al. The dynamic process of β(2)-adrenergic receptor activation. Cell 152, 532–542 (2013).

12. Gregorio, G. G. et al. Single-molecule analysis of ligand efficacy in β2AR-G-protein activation. Nature 547, 68–73 (2017).

13. Kobayashi, K. et al. The dynamic basis of G-protein recognition and activation by a GPCR. Nature 652, 812–821 (2026).

14. Vo, A. A. et al. Snapshots of the dynamic basis of NTSR1 G protein subtype promiscuity. Nature 652, 803–811 (2026).

15. Khan, S. et al. Structural snapshots capture nucleotide release at the μ-opioid receptor. Nature 648, 755–763 (2025).

16. Robertson, M. J. et al. Non-equilibrium snapshots of ligand efficacy at the μ-opioid receptor. Nature 652, 794–802 (2026).

17. Kato, H. E. et al. Conformational transitions of a neurotensin receptor 1-Gi1 complex. Nature 572, 80–85 (2019).

18. Kim, T. H. et al. The role of ligands on the equilibria between functional states of a G protein-coupled receptor. J. Am. Chem. Soc. 135, 9465–9474 (2013).

19. Ye, L., Van Eps, N., Zimmer, M., Ernst, O. P. & Prosser, R. S. Activation of the A2A adenosine G-protein-coupled receptor by conformational selection. Nature 533, 265–268 (2016).

20. Huang, S. K. et al. Delineating the conformational landscape of the adenosine A2A receptor during G protein coupling. Cell 184, 1884–1894.e14 (2021).

21. Hietpas, R. T., Jensen, J. D. & Bolon, D. N. A. Experimental illumination of a fitness landscape. Proc. Natl. Acad. Sci. U. S. A. 108, 7896–7901 (2011).

22. McLaughlin, R. N., Jr, Poelwijk, F. J., Raman, A., Gosal, W. S. & Ranganathan, R. The spatial architecture of protein function and adaptation. Nature 491, 138–142 (2012).

23. Faure, A. J. et al. Mapping the energetic and allosteric landscapes of protein binding domains. Nature 604, 175–183 (2022).

24. Leander, M., Liu, Z., Cui, Q. & Raman, S. Deep mutational scanning and machine learning reveal structural and molecular rules governing allosteric hotspots in homologous proteins. Elife 11, (2022).

25. Fowler, D. M. & Fields, S. Deep mutational scanning: a new style of protein science. Nat. Methods 11, 801–807 (2014).

26. Howard, M. K. & Coyote-Maestas, W. Mechanistic mutational scanning to uncover the secret life of proteins. Annu. Rev. Biomed. Data Sci. 9, 265–283 (2026).

27. Oakes, B. L. et al. Profiling of engineering hotspots identifies an allosteric CRISPR-Cas9 switch. Nat. Biotechnol. 34, 646–651 (2016).

28. Matreyek, K. A. et al. Multiplex assessment of protein variant abundance by massively parallel sequencing. Nat. Genet. 50, 874–882 (2018).

29. Starita, L. M. et al. Massively parallel functional analysis of BRCA1 RING domain variants. Genetics 200, 413–422 (2015).

30. Coyote-Maestas, W. et al. Probing ion channel functional architecture and domain recombination compatibility by massively parallel domain insertion profiling. Nat. Commun. 12, 7114 (2021).

31. Yee, S. W. et al. The full spectrum of SLC22 OCT1 mutations illuminates the bridge between drug transporter biophysics and pharmacogenomics. Mol. Cell 84, 1932–1947.e10 (2024).

32. Howard, M. K. et al. Molecular basis of proton sensing by G protein-coupled receptors. Cell 188, 671–687.e20 (2025).

33. Beltran, A., Naqvi, M. M., Faure, A. J. & Lehner, B. The allosteric landscape of the Src kinase. Sci. Adv. 12, eaea2726 (2026).

34. McCormick, J. W., Russo, M. A., Thompson, S., Blevins, A. & Reynolds, K. A. Structurally distributed surface sites tune allosteric regulation. Elife 10, e68346 (2021).

35. Coyote-Maestas, W., Nedrud, D., He, Y. & Schmidt, D. Determinants of trafficking, conduction, and disease within a K+ channel revealed through multiparametric deep mutational scanning. Elife 11, e76903 (2022).

36. Heydenreich, F. M. et al. Molecular determinants of ligand efficacy and potency in GPCR signaling. Science 382, eadh1859 (2023).

37. Jones, E. M. et al. Structural and functional characterization of G protein-coupled receptors with deep mutational scanning. Elife 9, e54895 (2020).

38. Howard, C. J. et al. High-resolution deep mutational scanning of the melanocortin-4 receptor enables target characterization for drug discovery. Elife 13, RP104725 (2025).

39. Che, T. & Roth, B. L. Molecular basis of opioid receptor signaling. Cell 186, 5203–5219 (2023).

40. Ramos-Gonzalez, N., Paul, B. & Majumdar, S. IUPHAR themed review: Opioid efficacy, bias, and selectivity. Pharmacol. Res. 197, 106961 (2023).

41. Chan, J. S., Chiu, T. T. & Wong, Y. H. Activation of type II adenylyl cyclase by the cloned mu-opioid receptor: coupling to multiple G proteins. J. Neurochem. 65, 2682–2689 (1995).

42. Ueda, H. et al. Reconstitution of rat brain mu opioid receptors with purified guanine nucleotide-binding regulatory proteins, Gi and Go. Proc. Natl. Acad. Sci. U. S. A. 85, 7013–7017 (1988).

43. Olsen, R. H. J. et al. TRUPATH, an open-source biosensor platform for interrogating the GPCR transducerome. Nat. Chem. Biol. 16, 841–849 (2020).

44. Qu, Q. et al. Insights into distinct signaling profiles of the µOR activated by diverse agonists. Nat. Chem. Biol. 19, 423–430 (2023).

45. Macdonald, C. B. et al. DIMPLE: deep insertion, deletion, and missense mutation libraries for exploring protein variation in evolution, disease, and biology. Genome Biol. 24, 36 (2023).

46. Macdonald, C., et al. DIMPLE library generation and assembly protocol. https://www.protocols.io/view/dimple-library-generation-and-assembly-protocol-eb5fbaq3p (2025).

47. Matreyek, K. A., Stephany, J. J., Chiasson, M. A., Hasle, N. & Fowler, D. M. An improved platform for functional assessment of large protein libraries in mammalian cells. Nucleic Acids Res. 48, e1 (2020).

48. Koehl, A. et al. Structure of the µ-opioid receptor-Gi protein complex. Nature 558, 547–552 (2018).

49. Fenalti, G. et al. Molecular control of δ-opioid receptor signalling. Nature 506, 191–196 (2014).

50. Katritch, V. et al. Allosteric sodium in class A GPCR signaling. Trends Biochem. Sci. 39, 233–244 (2014).

51. Zhou, Q. et al. Common activation mechanism of class A GPCRs. Elife 8, e50279 (2019).

52. Ballesteros, J. A. & Weinstein, H. [19] Integrated methods for the construction of three-dimensional models and computational probing of structure-function relations in G protein-coupled receptors. in Methods in Neurosciences (ed. Sealfon, S. C.) vol. 25 366–428 (Elsevier, 1995).

53. Pert, C. B., Pasternak, G. & Snyder, S. H. Opiate agonists and antagonists discriminated by receptor binding in brain. Science 182, 1359–1361 (1973).

54. Pert, C. B. & Snyder, S. H. Opiate receptor binding of agonists and antagonists affected differentially by sodium. Mol. Pharmacol. 10, 868–879 (1974).

55. Surratt, C. K. et al. -mu opiate receptor. Charged transmembrane domain amino acids are critical for agonist recognition and intrinsic activity. J. Biol. Chem. 269, 20548–20553 (1994).

56. Manglik, A. et al. Structure-based discovery of opioid analgesics with reduced side effects. Nature 537, 185–190 (2016).

57. Schmid, C. L. et al. Bias factor and therapeutic window correlate to predict safer opioid analgesics. Cell 171, 1165–1175.e13 (2017).

58. Váradi, A. et al. Mitragynine/corynantheidine pseudoindoxyls as opioid analgesics with mu agonism and delta antagonism, which do not recruit β-arrestin-2. J. Med. Chem. 59, 8381–8397 (2016).

59. Faouzi, A. et al. Structure-based design of bitopic ligands for the µ-opioid receptor. Nature 613, 767–774 (2023).

60. Shulman, M., Wai, J. M. & Nunes, E. V. Buprenorphine treatment for opioid use disorder: An overview. CNS Drugs 33, 567–580 (2019).

61. Zhuang, Y. et al. Molecular recognition of morphine and fentanyl by the human μ-opioid receptor. Cell 185, 4361–4375.e19 (2022).

62. Chai Discovery, et al. Chai-1: Decoding the molecular interactions of life. *bioRxiv* (2024) doi:10.1101/2024.10.10.615955.

63. Konovalov, K., et al. Transition intermediates encode ligand efficacy at the μ-opioid receptor-GI protein complex. bioRxiv (2026) doi:10.64898/2026.09.02.748988.

64. Ravindranathan, A. et al. Functional characterization of human variants of the mu-opioid receptor gene. Proc. Natl. Acad. Sci. U. S. A. 106, 10811–10816 (2009).

65. Kofuku, Y. et al. Efficacy of the β₂-adrenergic receptor is determined by conformational equilibrium in the transmembrane region. Nat. Commun. 3, 1045 (2012).

66. Su, M. et al. Structures of β1-adrenergic receptor in complex with Gs and ligands of different efficacies. Nat. Commun. 13, 4095 (2022).

67. Cong, X. et al. Molecular insights into the biased signaling mechanism of the μ-opioid receptor. Mol. Cell 81, 4165–4175.e6 (2021).

68. Sounier, R. et al. Propagation of conformational changes during μ-opioid receptor activation. Nature 524, 375–378 (2015).

69. Ball, D. I. et al. Salmeterol, a novel, long-acting beta 2-adrenoceptor agonist: characterization of pharmacological activity in vitro and in vivo. Br. J. Pharmacol. 104, 665–671 (1991).

70. Burris, K. D. et al. Aripiprazole, a novel antipsychotic, is a high-affinity partial agonist at human dopamine D2 receptors. J. Pharmacol. Exp. Ther. 302, 381–389 (2002).

71. Davis, M. Buprenorphine pharmacodynamics: A bridge to understanding buprenorphine clinical benefits. Drugs 85, 215–230 (2025).

72. Dahan, A. et al. Buprenorphine induces ceiling in respiratory depression but not in analgesia. Br. J. Anaesth. 96, 627–632 (2006).

73. Yang, F. et al. Different conformational responses of the β2-adrenergic receptor-Gs complex upon binding of the partial agonist salbutamol or the full agonist isoprenaline. Natl. Sci. Rev. 8, nwaa284 (2021).

74. Masureel, M. et al. Structural insights into binding specificity, efficacy and bias of a β2AR partial agonist. Nat. Chem. Biol. 14, 1059–1066 (2018).

75. Rao, J., et al. Rosace: a robust deep mutational scanning analysis framework employing position and mean-variance shrinkage. *bioRxiv* 2023.10.24.562292 (2023) doi:10.1101/2023.10.24.562292.

76. Bushnell, B. BBTools software package. Preprint at (2014).

77. Van Der Auwera, G. A. & O’Connor, B. D. Genomics in the Cloud: Using Docker, Gatk, and Wdl in Terra. (O’Reilly Media, 2020).

78. Freudenberg, J. et al. Accurate variant effect estimation in FACS-based deep mutational scanning data with Lilace. Genome Biol. 27, 48 (2026).

79. DiBerto, J. F., Smart, K., Olsen, R. H. J. & Roth, B. L. Agonist and antagonist TRUPATH assays for G protein-coupled receptors. STAR Protoc. 3, 101259 (2022).

80. Staus, D. P. et al. Sortase ligation enables homogeneous GPCR phosphorylation to reveal diversity in β-arrestin coupling. Proc. Natl. Acad. Sci. U. S. A. 115, 3834–3839 (2018).

81. Maeda, S. et al. Development of an antibody fragment that stabilizes GPCR/G-protein complexes. Nat. Commun. 9, 3712 (2018).

82. CryoSPARC Team et al. End-to-end automation of repeat-target cryo-EM structure determination in CryoSPARC. *bioRxiv* 2025.10.17.682689 (2025) doi:10.1101/2025.10.17.682689.

83. Kim, K., Li, H. & Clarke, O. High-resolution ab initio reconstruction enables cryo-EM structure determination of small particles. *bioRxiv* 2025.09.08.674935 (2025) doi:10.1101/2025.09.08.674935.

84. Pettersen, E. F. et al. UCSF ChimeraX: Structure visualization for researchers, educators, and developers. Protein Sci. 30, 70–82 (2021).

85. Liebschner, D. et al. Macromolecular structure determination using X-rays, neutrons and electrons: recent developments in Phenix. Acta Crystallogr. D Struct. Biol. 75, 861–877 (2019).

86. Emsley, P., Lohkamp, B., Scott, W. G. & Cowtan, K. Features and development of coot. Acta Crystallogr. D Biol. Crystallogr. 66, 486–501 (2010).

